# Oncogenic herpesvirus engages the endothelial transcription factors SOX18 and PROX1 to increase viral genome copies and virus production

**DOI:** 10.1101/742932

**Authors:** Silvia Gramolelli, Endrit Elbasani, Veijo Nurminen, Krista Tuohinto, Thomas Günther, Riikka E. Kallinen, Seppo P. Kaijalainen, Raquel Diaz, Adam Grundhoff, Caj Haglund, Joseph M. Ziegelbauer, Mark Bower, Mathias Francois, Päivi M. Ojala

**Affiliations:** Translational Cancer Medicine Research Program, Faculty of Medicine, University of Helsinki, Helsinki, 00014; Finland; Heinrich Pette Institute, Leibniz Institute for Experimental Virology, Hamburg, 205052; Germany; Department of Surgery, University of Helsinki and Helsinki University Hospital, Helsinki, 00014; Finland; HIV and AIDS Malignancy Branch, Center for Cancer Research, National Cancer Institute, National Institutes of Health, Bethesda, MD 20892; USA; National Centre for HIV Malignancy, Chelsea & Westminster Hospital, London, SW10 9NH; UK; Institute of Molecular Biosciences, The University of Queensland, Brisbane, QLD 4072; Australia; Department of Infectious Diseases, Imperial College London, London, W2 1PG; United Kingdom

**Keywords:** Kaposi sarcoma, KSHV, herpesvirus, K8.1, transcription factors, PROX1, SOX18, lymphatic endothelial cells, viral genome replication, viral reactivation

## Abstract

Kaposi sarcoma (KS) is a tumour of endothelial origin caused by KS herpesvirus (KSHV) infection and suggested to originate from lymphatic endothelial cells (LECs). While KSHV establishes latency in virtually all susceptible cell types, LECs support a spontaneous lytic gene expression program with high viral genome copies and release of infectious virus. Here, we investigated the role of PROX1, SOX18 and COUPTF2, drivers of lymphatic endothelial fate during embryogenesis, in this unique KSHV infection program. We found that these factors were co-expressed in KS tumours with the viral lytic marker K8.1, and that SOX18 and PROX1 regulate KSHV infection via two independent mechanisms. SOX18 binds to the viral origins of replication and its depletion or chemical inhibition significantly reduced the KSHV genome copies in LECs. PROX1 interacts with ORF50, the initiator of the lytic cascade, increases lytic gene expression and virus production and its depletion reduces KSHV spontaneous lytic reactivation. Upon lytic replication, PROX1 binds to the KSHV genome in the promoter region of ORF50 and enhances its transactivation activity. These results demonstrate the importance of two endothelial transcription factors in the regulation of the KSHV life cycle and introduce SOX18 inhibition as a potential, novel therapeutic modality for KS.

## MAIN

Kaposi sarcoma (KS) is an angiogenic endothelial tumour caused by KS herpesvirus (KSHV). KS is the most common cancer among AIDS patients and a frequent malignancy in Sub-Saharan Africa^1^. KS presents with skin lesions that progress from patch to plaque and ultimately to nodular tumours; in advanced disease visceral involvement is also seen^2, 3^. The histopathological hallmark of KS is the presence of KSHV-positive spindle cells (SC), the tumour cells of KS^2, 3^. The cell of origin of SC has been debated for two decades^3^. The prevailing hypothesis suggests lymphatic endothelial origin, although blood endothelial cells or mesenchymal cells are also candidates^1, 4^. *In vitro*, latency is the default replication program in KSHV-infected cells with undetectable levels of lytic genes expressed. However, KSHV infection of lymphatic, but not blood, endothelial cells (LEC and BEC) leads to a unique infection program characterized by high KSHV genome copies, spontaneous lytic gene expression and release of infectious virus^5–8^.

During embryonic development, LEC precursors originate from COUPTF2/SOX18 double-positive BECs that physically separate from the cardinal vein to establish a primary lymphatic vascular plexus. In this process, COUPTF2 and SOX18 drive the expression of *PROX1* thereby orchestrating LEC differentiation^9, 10^. Here, we explored the role of these transcription factors (TF), instrumental for establishment and maintenance of LEC identity, on the spontaneously lytic KSHV infection program in LEC.

We identified SOX18 and PROX1, both expressed in KSHV-positive SCs in tumours, as key regulators of the unique KSHV infection program in LECs. SOX18 binds to the viral origins of replication and increases viral genome copies. Chemical inhibition or depletion of SOX18 reduced intracellular viral genomes and titers. PROX1 enhances the expression of KSHV lytic genes and virus production. During the lytic cycle, it binds to the promoter region of *ORF50*, the viral initiator of the lytic cascade. This study uncovers that TFs essential for maintaining the LEC identity are utilized by KSHV to support its spontaneous lytic infection program in LECs, the proposed cells of origin of KS-SCs.

## RESULTS

### PROX1, SOX18, COUPTF2 and K8.1 are expressed in KS tumours

To study the expression of PROX1, SOX18 and COUPTF2 in KS tumours, we stained consecutive sections from nine KS patients (seven AIDS-KS and two classic-KS) and compared their expression to skin from KS-negative donors. While PROX1 expression in KS tumours has been shown previously^11–14^, SOX18 and COUPTF2 expression has not been reported. Consecutive KS sections were also stained for latency-associated nuclear antigen (LANA) and K8.1, markers of KSHV latent and lytic infection, respectively. In the analysed KS tumours, SOX18, PROX1 and COUPTF2 were highly expressed while in KS-negative skin samples their expression was restricted to the vascular endothelium, as expected. Notably, SOX18, PROX1 and COUPTF2, all followed the distribution of LANA and K8.1, indicating that these TFs are expressed in SCs together with the late lytic protein K8.1 (Fig.1a,b, Supplementary Fig.1a,b). K8.1 and other lytic transcripts have been recently detected in KS lesions by RNA-Seq^15^. Here, we observed a cytoplasmic K8.1 staining, consistent with the predicted distribution of the protein. No K8.1 signal was detected on either normal skin or using an isotype control antibody on KS sections, confirming the specificity of the staining (Supplementary Fig.1c, d).

**Fig. 1.**
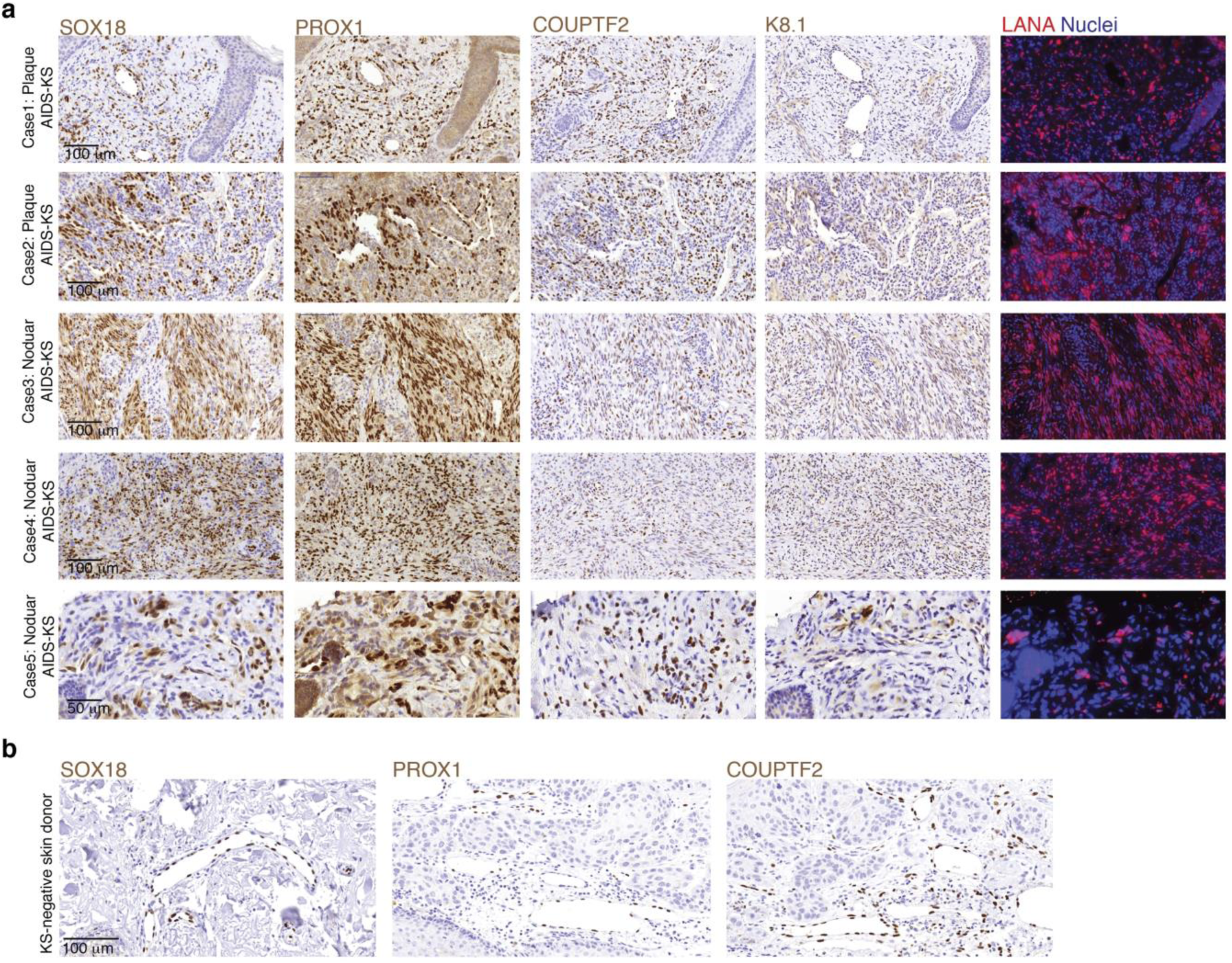
PROX1, SOX18, COUPTF2 and K8.1 are expressed in KS tumours. (a,b) Representative images of consecutive sections from (a) five AIDS-KS biopsies and (b) Skin from a KS negative donor stained with the indicated antibodies.

Overall, SOX18, PROX1 and COUPTF2 are expressed in KS-SC together with latent (LANA) and late lytic (K8.1) proteins, supporting the relevance to study their role in the KSHV life cycle.

### Endothelial TFs and lytic genes are highly expressed in KSHV-infected LECs

KSHV infection has been shown to skew the LEC and BEC identity and modulate PROX1 transcription so that it is downregulated in KSHV-infected LECs (KLECs) and upregulated in KSHV-infected BECs (KBECs)^12, 14, 16–18^. In these studies, however, SOX18 and COUPTF2 levels were not analysed and PROX1 protein levels were not rigorously compared. To address this, BECs and LECs were infected with rKSHV.219, a recombinant virus expressing GFP and RFP during latent and lytic phase, respectively^19^.

The expression levels of PROX1, SOX18 and COUPTF2, together with infection (GFP), latent (LANA) and lytic (ORF50, ORF45 and K8.1) markers were monitored either during a 14 day-time course of infection or analysed 14 days post-infection (d.p.i.) (Fig.2a-e, Supplementary Fig.2 a-c). As previously reported^12, 14^, *PROX1* transcripts, initially low in BEC, were upregulated in KBECs increasing up to three-fold by 14.d.p.i. and downregulated to four-fold in KLECs by 14 d.p.i. (Fig.2a). KSHV infection increased *SOX18* transcripts in KBECs (three-fold by 10 d.p.i.) and KLECs (three-fold already at 3 d.p.i.), while *COUPTF2* expression did not change (Fig.2a). While we observed an initial expression of lytic markers in KBECs, followed by their downregulation and latency establishment, KSHV gene expression program was different in KLECs. In accordance with the unique infection program in KLECs, the lytic expression increased throughout the timepoints analysed, except at day10 p.i. where we consistently observed a downregulation of both viral and TFs proteins (Fig.2b,c,e, Supplementary Fig.2a).

**Fig. 2.**
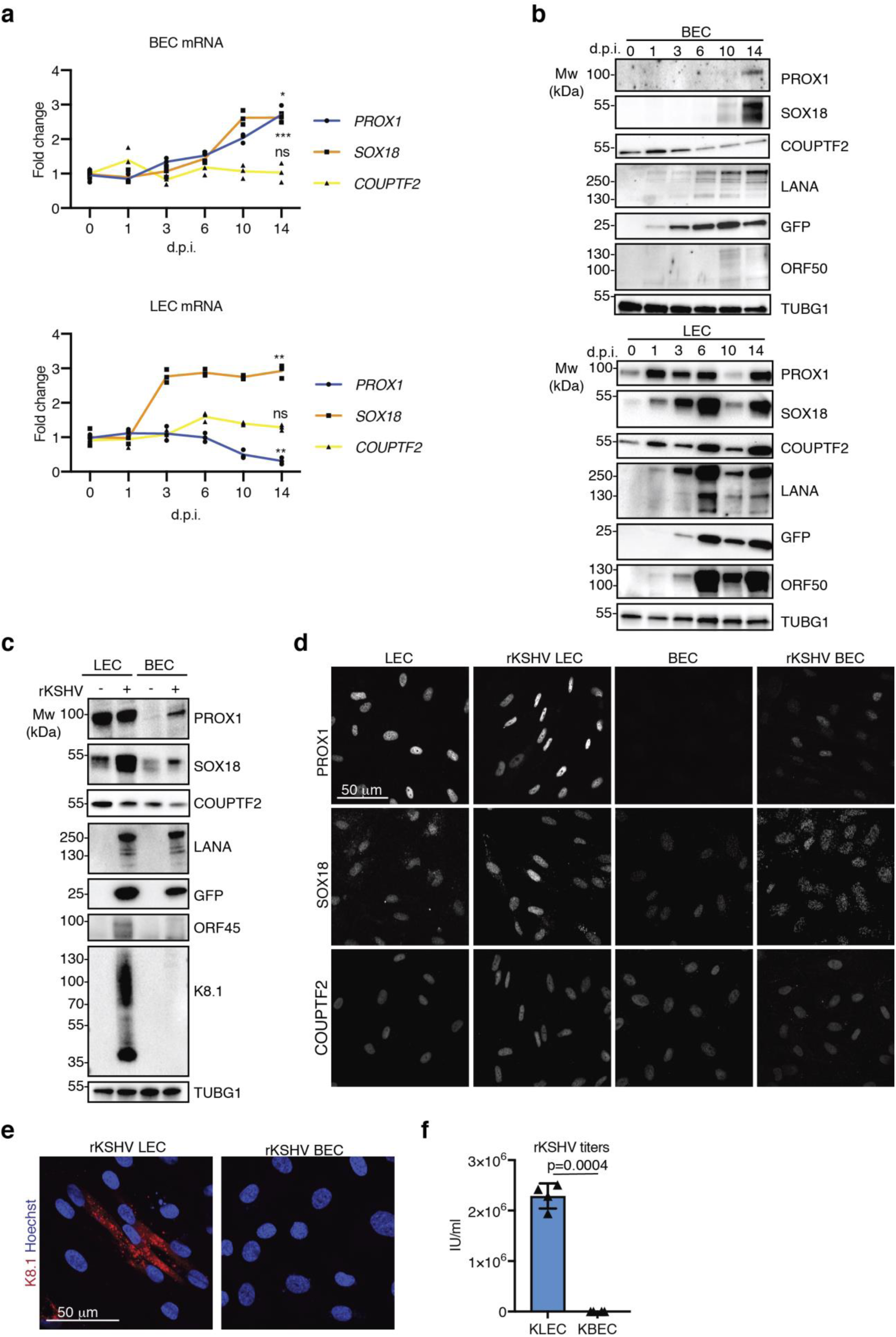
Endothelial TFs and lytic genes are highly expressed in KSHV-infected LECs. **(a-f)** Primary BECs and LECs were infected with rKSHV.219 or left uninfected and analysed at the indicated timepoints (a,b) and 14 d.p.i. (days post-infection) (c-f). (a) RT-qPCR for the indicated cellular targets at the indicated timepoint in BEC (upper panel) and LEC (lower panel); single data from n=3 independent experiments ± SD are shown for each timepoints; p values were calculated using ordinary one way-anova followed by Dunnett’s correction for multiple comparisons. *: p<0.033; **: p<0.02, ***: p<0.001. Exact p values are shown in Supplementary Table3. (b) Immunoblot of BEC (upper panel) and LEC (lower panel) treated as in (a) for the indicated targets and gamma-tubulin (TUBG1) as a loading control. The experiment was repeated three independent times. Representative cropped blots are shown, uncropped blots are shown in Supplementary Fig.6. (c) Immunoblot for the indicated cellular targets 14 d.p.i., and gamma-tubulin (TUBG1) as a loading control. The experiment was repeated three independent times. Representative cropped blots are shown, uncropped blots are shown in Supplementary Fig.6. (d) IF staining for PROX1, SOX18, and COUPTF2 in the indicated cell types treated as in (c). Representative images from two independent experiments are shown. (e) IF staining in the indicated cell types 14 d.p.i. for K8.1 (red), nuclei are counterstained with Hoechst 33342. Representative images from two independent experiments are shown. (**f)** Titration of cell free supernatant from KBECs and KLECs 14 d.p.i. Single values from n=4 biological replicates are shown. Bars represent mean ± SD. P value was calculated using two-tailed paired t-test.

Despite the transcriptional downregulation, comparable levels of PROX1 protein were detected by immunoblotting in LECs and KLECs infected with either rKSHV.219 or wild-type-(WT) KSHV and confirmed by immunofluorescence at single cell level (Fig.2b-e Supplementary Fig.2b,c). SOX18 protein levels were increased in both KLECs and KBECs, while COUPTF2 levels did not vary significantly (Fig.2b,c,d, Supplementary Fig.2b,c). At 14 d.p.i., spontaneous production of high titers (10^5^ - 10^6^ IU/ml) was observed only from KLECs, and not from KBECs infected with two different KSHV strains, rKSHV.219 and WT-KSHV (Fig.2f, Supplementary Fig.2d).

Thus, SOX18, PROX1 and COUPTF2 were all highly expressed in KLECs, which, compared to the latently-infected KBECs, expressed lytic markers and produced high titers. This points to KLECs as a physiologically-relevant model to study SOX18, PROX1 and COUPTF2 in KSHV infection.

### SOX18 and PROX1 regulate KSHV infection through different mechanisms

To investigate SOX18, PROX1 and COUPTF2 in KSHV infection, KLECs treated with control siRNA or siRNAs targeting these TFs for 72h were analysed for viral mRNAs, protein expression and titers (Fig.3a-c, Supplementary Fig.3a-d). *PROX1* and *SOX18* depletion significantly decreased the lytic transcript and protein levels, while *COUPTF2* depletion caused a modest increase in lytic gene expression. Of note, SOX18 depletion had a marked negative effect on *LANA* mRNA and protein levels while *PROX1* depletion reduced *LANA* transcripts to a half, but only marginally its protein levels (Fig.3a,b, Supplementary Fig.2a). KSHV titers were reduced by a half upon *PROX1* depletion and more severely after SOX18 silencing, while *COUPTF2* silencing did not affect the titers, despite the decreased lytic protein expression.

**Fig. 3.**
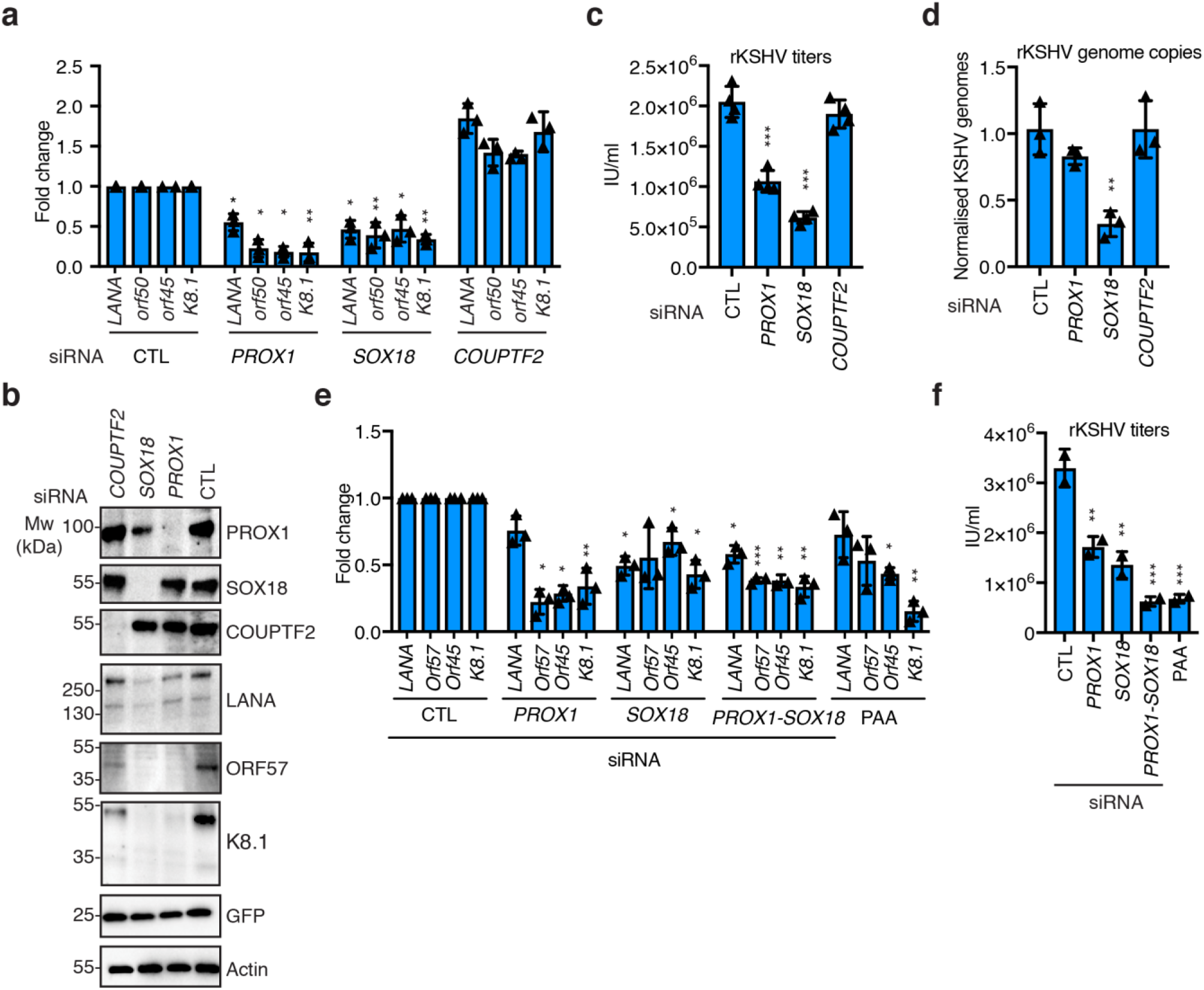
SOX18 and PROX1 regulate KSHV production through different mechanisms. **(a-d)** KLECs were treated 14 d.p.i. with the indicated siRNA for 72 h and analysed. (a) RT-qPCR for the indicated viral-encoded targets. Single values from n=3 independent experiments are shown. Bars represent mean ± SD. (b) Immunoblot for the indicated targets and actin as a loading control. The experiment was repeated three independent times. Representative cropped blots are shown, uncropped blots are shown in Supplementary Fig.6. (c) Titration for released infectious virus. Single values from n=3 independent replicates are shown. Bars represent mean ± SD (d) Relative number of intracellular KSHV genomes. Single values from n=3 independent experiments are shown. Bars represent mean ± SD. **(e,f)** KLECs were treated 14 d.p.i. with the indicated siRNAs or 0.5mM PAA for 72h. (e) Cells were analysed by RT-qPCR for the indicated viral targets. Single values from n=3 independent experiments are shown. Bars represent mean ± SD. (f) Titration of the cell-free supernatant. Single values from n=2 independent experiments are shown. Bars represent mean ± SD. P values were calculated in panels (a), (c-f) using ordinary one way-anova followed by Dunnett’s correction for multiple comparisons. *: p<0.033; **: p<0.02, ***: p<0.001. Exact p values are shown in Supplementary Table3.

In KLECs *PROX1* depletion did not change SOX18 levels whereas *SOX18* depletion decreased the *PROX1* mRNA and slightly its protein levels (Fig.3b, Supplementary Fig.3d). Since SOX18 positively regulates *PROX1* during LEC development, we explored the reciprocal effect of depletion of these TFs also in non-infected LECs. In the absence of KSHV, silencing of either *PROX1, SOX18* or *COUPTF2* negatively affected the levels of all the other TFs. This was particularly evident for PROX1 and even more for SOX18. PROX1 expression decreased upon both SOX18 and COUPTF2 silencing, and similarly, decreased SOX18 expression upon PROX1 and COUPTF2 depletion was observed in non-infected LECs (Supplementary Fig.3e). This suggests that in KLECs KSHV infection has uncoupled the reciprocal regulation of PROX1, SOX18 and COUPTF2 expression.

Since SOX18 depletion affected not only the lytic genes but also the latent LANA expression, we investigated whether this was due to a diminished number of intracellular viral genomes (episomes). While *PROX1* silencing modestly decreased viral episome numbers (by 10%), *SOX18* depletion decreased them by about 70% (Fig.3d), suggesting that SOX18 may contribute to the high episome copies in KLECs.

Next, we co-silenced *PROX1* and *SOX18* to test if they reduce KSHV lytic gene expression and titers independently of each other. As a positive control, we also treated KLECs with phosphonoacetic acid (PAA), an inhibitor of the viral DNA polymerase which inhibits the late lytic program and virus production (Fig.3e,f). As expected, PAA treatment decreased slightly transcription of the intermediate-early *ORF45* gene and more dramatically the late lytic *K8.1*, whereas *PROX1* and *SOX18* co-silencing reduced both viral gene expression (early and late) and the reduction in titers, more efficient if compared to their individual silencing, was comparable to PAA treatment.

These results suggest that PROX1 and SOX18 regulate the spontaneous virus production in KLECs via different mechanisms, SOX18 by controlling the KSHV episome numbers while PROX1 regulates lytic viral gene expression.

### SOX18 inhibition efficiently decreases KSHV genome copies and infectious virus release

SOX18 homo- and heterodimerization can be inhibited by a small molecule, SM4^20–22^ and by an FDA-approved beta blocker, propranolol^23^. Propranolol is produced as a 1:1 racemic mixture of R(+) and S(-) enantiomers. The S(-) form exhibits beta-adrenergic blocking activity, while the less potent R(+) enantiomer is carried over during the production. Similar to SM4, R(+) propranolol blocks SOX18 homo- and heterodimerization^23^, as a result, SOX18 fails to regulate a subset of its target genes^21^.

Given the significant reduction in viral episome numbers and titers upon *SOX18* silencing in KLECs, we explored whether its targeting by SM4 and R(+) propranolol would produce a similar outcome. KLECs were treated with SM4, the propranolol racemic mixture (R+S) or the pure enantiomers R(+) and S(-) (Fig.4a-d) using the same range of inhibitor concentrations shown to inhibit SOX18-mediated activation of EC-specific genes^21^. We observed a dose-dependent decrease in KSHV episomes and titers in SM4 and R(+)propranolol-treated KLECs compared to the DMSO-treated control (Fig.4a-d). The racemic propranolol and the S(-) enantiomer increased both KSHV episomes and titers. This agrees with a previous report where propranolol treatment of immortalized KLECs induced to lytic cycle by phorbol esters increased lytic KSHV gene expression and titers, likely by interfering with the cell cycle^24^, rather than having a SOX18-specific effect.

**Fig. 4.**
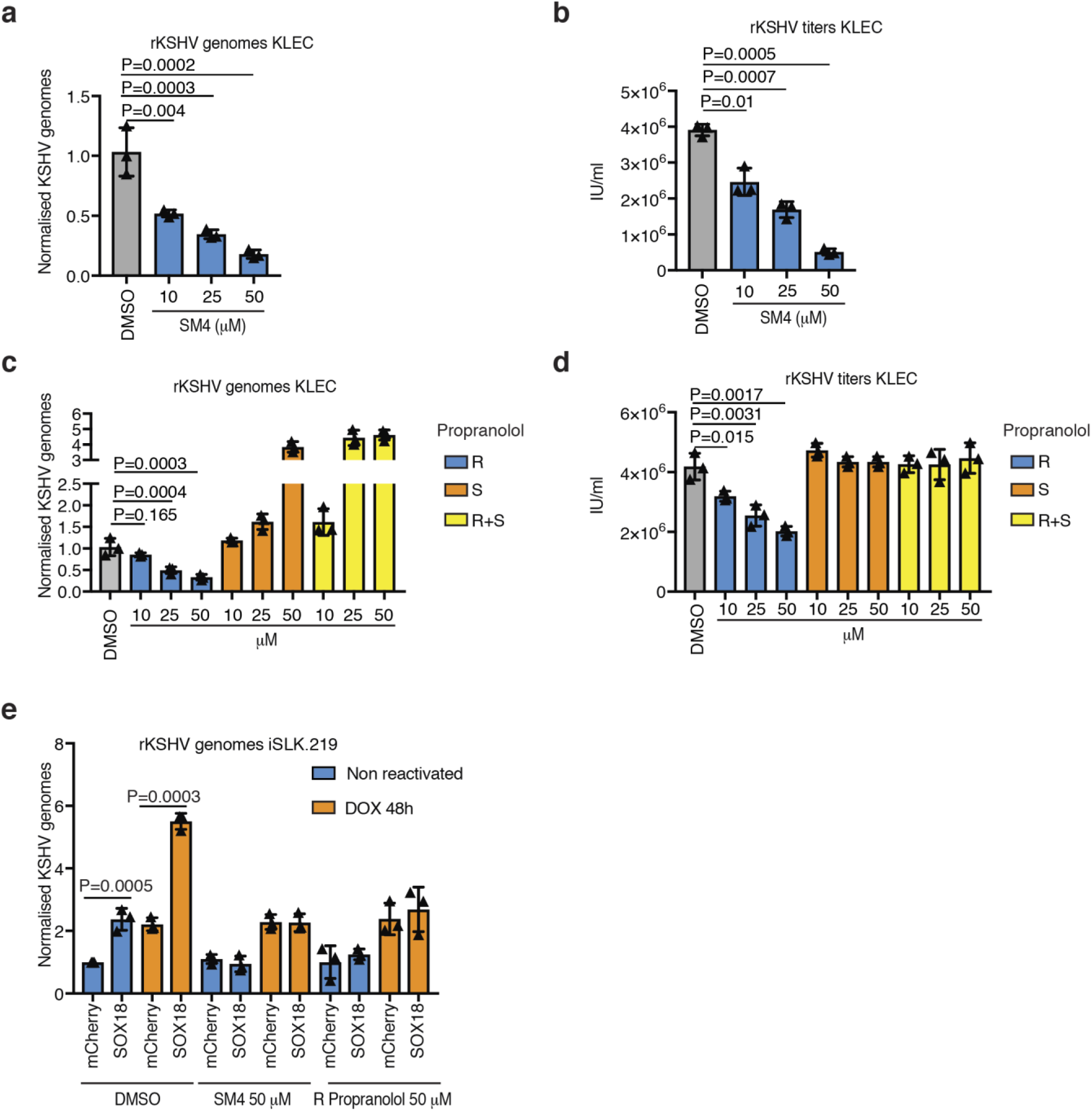
Selective inhibition of SOX18 efficiently decreases the number of KSHV episomes. **(a-d)** KLECs 10 d.p.i. were treated for six days with the indicated compounds at the concentrations shown. The relative number of intracellular KSHV genome copies was quantified (a,c), and the released infectious virus in the supernatant was measured (b,d). Single values from n=3 independent replicates are shown. Bars represent mean ± SD. P values were calculated using ordinary one way-anova followed by Dunnett’s correction for multiple comparisons. Exact p values are shown. **(e)** iSLK.219 were transduced with lentiviruses expressing either SOX18 or NLS-mCherry (indicated as mCherry) vector control. One day later cells were treated either with DMSO or with the indicated drug for 48h in the presence (orange bars) or absence (blue bars) of dox for the induction of the lytic cycle. The relative number of intracellular KSHV genome copies was quantified. Single values from n=3 biological experiments are shown. Bars represent mean ± SD. P value were calculated using two-tailed paired t-test.

To further assess the specificity of the inhibitors, we tested these drugs on the SOX18-negative iSLK.219 cells transduced with a control (myc-NLS-mCherry) or a myc-tagged SOX18-expressing lentivirus. iSLK.219 is a cancer-derived cell line stably infected with rKSHV.219 and expressing KSHV-*ORF50* that drives lytic reactivation from a doxycycline (dox)-inducible promoter^25^. In the absence of dox these cells are strictly latent. KSHV episomes were quantified in latently-infected cells or cells reactivated with dox for 48h and treated with the indicated inhibitors or DMSO (Fig.4e). SOX18 expression significantly increased KSHV episome copies both in latent and lytic cells. SM4 and R(+) propranolol treatments reduced episome copies in iSLK.219 ectopically expressing SOX18 but not in the controls, supporting the specificity of the SOX18 chemical inhibition.

Therefore, these results provide a proof of principle that targeting SOX18 could represent a future strategy for KS treatment.

### SOX18 binds to the KSHV genome in the proximity of the origins of replication

To further investigate the effect of SOX18 on the KSHV episomes, iSLK.219 cells were transduced with the mCherry- or increasing amounts of SOX18-expressing lentiviruses and KSHV episomes were quantified 48h post-transduction (Fig.5a). We observed a significant, dose-dependent increase of KSHV episome copy number. We tested whether this was due to increased proliferation of SOX18-expressing iSLK.219 by 5-ethynyl-2’-deoxyuridine (EdU) treatment (Supplementary Fig.5a). No change in the percentage of proliferating (EdU-positive) cells was found following SOX18 expression. Additionally, KSHV gene, protein expression and titers were monitored for 72h in iSLK.219 after dox-induced reactivation (Supplementary Fig.5b-d). SOX18 ectopic expression mildly increased KSHV lytic transcripts and proteins without increasing the titers. These data suggest that SOX18 expression increases viral genome copies without increasing cell proliferation and only slightly contributes to lytic gene expression upon dox induction.

**Fig. 5.**
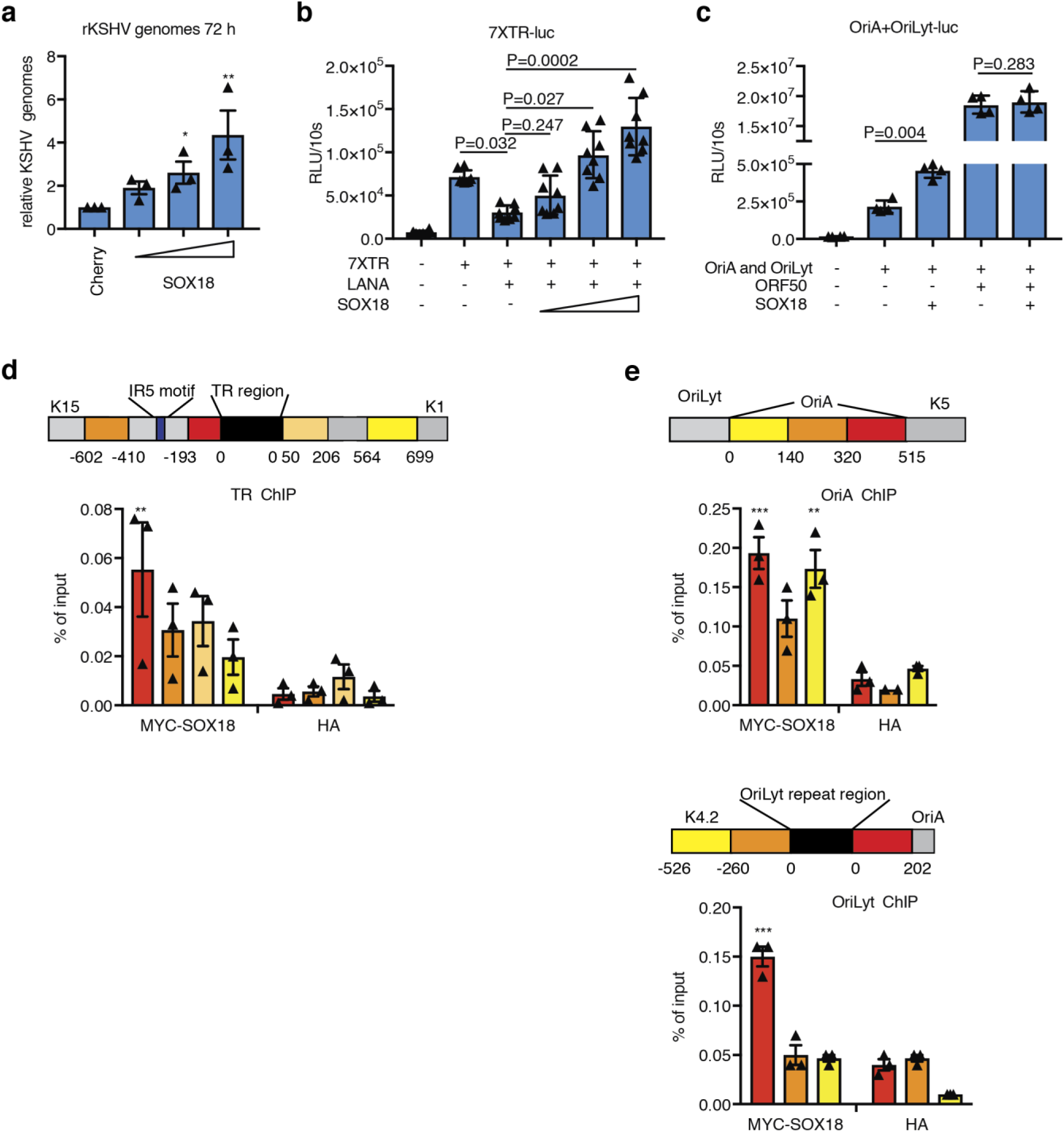
SOX18 binds to the proximity of KSHV origins of replication. **(a)** iSLK.219 were transduced with increasing amounts of lentiviruses expressing either SOX18 or mCherry vector control. 48h later, the relative number of intracellular KSHV genome copies was quantified. Single values for n=3 biological replicates are shown. Bars represent mean ± SD. **(b,c)** Luciferase reporter assays in HeLa cells transfected as indicated. Single values from (b) n=8 and (c) n=4 biological replicates are shown. Bars represent mean ± SEM. **(d;e)** Upper panels: schematic representation of the promoter regions amplified by qPCR following chromatin immunoprecipitation (ChIP) using either myc antibody to precipitate myc-tagged SOX18 or control HA antibody; numbers indicate the nucleotides upstream or downstream of the black regions; bottom panels: ChIP in iSLK.219 transduced with lentivirus expressing Myc-tagged SOX18. Single values for n=3 biological replicates are shown. Bars represent mean ± SEM. P values in all panels were calculated using ordinary one way-anova followed by Dunnett’s correction for multiple comparisons; *: p<0.033; **: p<0.02, ***: p<0.001. Exact p values are shown in Supplementary Table3.

During latency, KSHV replication occurs mainly by a LANA-dependent mechanism from the terminal-repeat (TR) region of the KSHV genome and, to a minor extent, through a LANA-independent mechanism from the OriA region, adjacent to the origin of lytic replication (OriLyt)^26, 27^. We measured the activity of luciferase-reporters harbouring either seven copies of the TR region (7XTR) or the OriA fused to OriLyt upstream of an SV40 promoter and a firefly luciferase reporter (Fig.5b,c, Supplementary Fig.5e,f). ORF50, that binds to the OriLyt and is a potent activator of this reporter^28^, was used as a positive control. In the presence of LANA, SOX18 expression increased the activity of the 7XTR reporter in a dose-dependent manner. SOX18 expression also increased the activity of the OriA+OriLyt reporter in an ORF50-independent manner. SOX18 did not change the activity of a reporter plasmid harbouring the *ORF5*0 promoter (Supplementary Fig.5g,) supporting the specificity of the activation observed in the 7XTR and OriA+OriLyt reporters.

Since the reporter assays suggested that SOX18 might bind to the KSHV origins of replication, we assessed this further by ChIP-qPCR in iSLK.219 ectopically expressing a myc-tagged SOX18 (Fig.5d,e). Binding of SOX18 to the KSHV genome was observed in regions adjacent to the TR and within and adjacent to the OriA region. SOX18 drives EC fate through the regulation of a subset of genes harbouring in their promoter an IR5 consensus motif, composed of two SOX18-binding sites spaced by five nucleotides (nt)^21^. Notably, one of these motifs was located adjacent to the KSHV-TR region bound by SOX18 (Fig.5d).

Overall these results indicate that SOX18 binds at the proximity of KSHV origins of latent replication, and it increases in viral episome copies.

### PROX1 enhances viral gene expression, binds to ORF50 and its promoter during KSHV lytic replication cycle

To obtain a comprehensive picture of the PROX1-mediated KSHV lytic gene regulation, we performed RNA-seq of KLECs transfected for 72 h with either control or PROX1-targeting siRNA and of dox-treated (24h) iSLK.219 cells expressing a myc-tagged PROX1 from a lentivirus. PROX1 silencing in KLECs significantly reduced expression of viral genes and viral circular RNAs, expressed during the lytic cycle^29, 30^ (Fig.6a, Supplementary Fig.5a). Analysis of differentially expressed genes (DEG; Supplementary Table1) revealed several cellular pathways involved in oncogenesis (e.g. p53 signalling, viral carcinogenesis, and transcription misregulation in cancer) altered by PROX1 silencing in KLECs (Supplementary Fig.5b), supporting the pivotal role of PROX1 in KS pathogenesis. Conversely, PROX1 reintroduction in iSLK.219 cells induced with dox led to an overall increase in viral gene expression (Fig.6a). The analysis of DEG (Supplementary Table2) in iSLK.219 did not identify differentially regulated pathways.

**Fig. 6.**
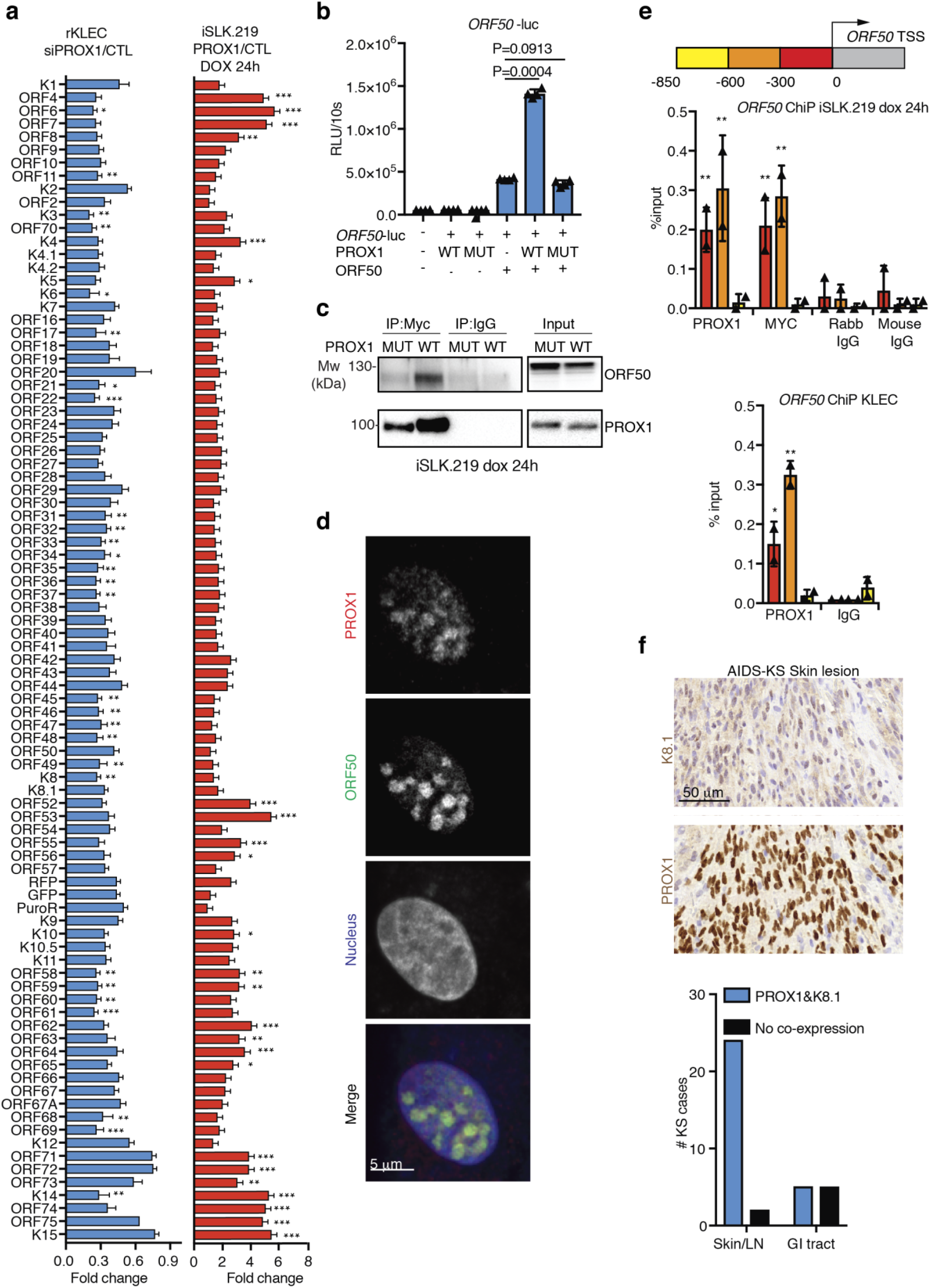
PROX1 is a positive regulator of viral gene expression, binds to ORF50 and to *ORF50* promoter during KSHV lytic replication cycle. **(a)** KLECs (left) and iSLK.219 (right) were treated in n=3 independent replicates, as indicated and subjected to RNA-seq. Fold change ±SEM in the viral transcripts over the appropriate control is shown. **(b)** Luciferase reporter assay in HEK293FT transfected with the indicated plamids for 36h. Single values for n=4 biological replicates are shown, bars represent mean ± SD. **(c)** Co-Immunoprecipitation of Myc-tagged PROX1 WT or MUT with ORF50 in iSLK.219 cells reactivated (dox) for 24h. The experiment was repeated two independent times. Representative cropped blots are shown, uncropped blots are shown in Supplementary Fig.6. **(d)** Immunofluorescence for PROX1 (red) and ORF50 (green) in KLECs infected with WT KSHV 16 d.p.i. A representative image is shown, the experiment was done two times. **(e)** Upper panel: schematic representation of the *ORF50* promoter regions amplified by qPCR following Chromatin immunoprecipitation (ChIP); numbers indicate the nucleotides upstream of the *ORF50* TSS; middle panel: ChIP in iSLK.219 transduced with lentivirus expressing Myc-tagged PROX1 reactivated (dox) for 24 h, chromatin was immunoprecipitated with two antibodies for PROX1 (anti-PROX1 and anti-Myc) and DNA was amplified in the promoter regions of the *ORF50* gene; lower panel: ChIP in KLECs 14 d.p.i. using PROX1 antibody as above. Bars represent mean ± SD. (**f)** KS lesions from skin or lymph node (LN) lesions from 26 patients and gastro-intestinal (GI) KS lesions from 10 patients were analysed for co-distribution of PROX1 and K8.1 staining in consecutive sections (left panel); representative image (right panels). P values in panels a,b,e were calculated using ordinary one way-anova followed by Dunnett’s correction for multiple comparisons. *: p<0.033; **: p<0.02, ***: p<0.001. Exact p values are shown in Supplementary Table3.

We further characterized iSLK.219 cells transduced with lentiviruses encoding a Myc-tagged PROX1, either WT or a mutant (MUT) harbouring two point mutations at the DNA binding site (N624A and N626A) and lacking transcriptional activity^31^, as well as a control vector. One day after dox-induced lytic reactivation PROX1 WT, but not MUT, expression increased the transcription of lytic KSHV genes and immunoblot analysis revealed higher lytic protein levels in the presence of PROX1 WT (Supplementary Fig.5c,d). High-content image analysis in iSLK.219 cells 24h after dox-induced reactivation, revealed that PROX1 WT, but not MUT, increased the percentage of cells positive for early (RFP, ORF57) and late (K8.1) lytic markers (Supplementary Fig.5e). KSHV titers were significantly higher in iSLK.219 cells expressing PROX1 WT when compared to MUT or controls (Supplementary Fig.5f). These results suggest that, upon lytic reactivation, a transcriptionally active PROX1 increases KSHV lytic gene expression and titers.

To assess if PROX1 could induce spontaneous lytic reactivation in latent iSLK.219, they were transduced with PROX1 (either WT or MUT) lentiviruses or controls in the absence of dox. This led to about two- to ten-fold increase in lytic mRNAs, but not in lytic proteins (Supplementary Fig.5g, h). Therefore, we hypothesized that an early viral lytic factor was needed for PROX1 to significantly enhance KSHV lytic cycle. Since KSHV-ORF50 is sufficient to trigger the complete lytic cascade, we investigated whether PROX1 would act in concert with ORF50 to increase the lytic gene expression. We first performed a luciferase-reporter assay using the *ORF50* promoter upstream of a luciferase gene (*ORF50*-luc) and measured its activity upon ectopic expression of PROX1, WT or MUT, or ORF50 expression plasmid alone and in combination with the PROX1 constructs (Fig.6b, Supplementary Fig.4i). PROX1 alone did not increase the *ORF50* promoter activity beyond the basal levels, in agreement with its inability to induce a strong, spontaneous reactivation in iSLK.219 cells. However, PROX1 WT, but not the transcriptionally inactive MUT, enhanced significantly the *ORF50*-mediated transcription from its own promoter. We also found that PROX1 WT, but not PROX1 MUT, co-immunoprecipitated ORF50 in iSLK.219 reactivated for 24h by dox-treatment (Fig.6c). Furthermore, in LECs infected with WT-KSHV, ORF50 and PROX1 co-localized in the viral replication and transcription compartments^32, 33^ (Pearson’s correlation coefficient, PCC=0.65, Fig.6d). To test whether PROX1 would bind to the *ORF50* promoter in infected cells, ChIP was performed in iSLK.219 (latent or dox-induced for 24h) to precipitate the Myc-tagged PROX1-bound chromatin with two different antibodies (anti-PROX1 and anti-Myc) followed by RT-qPCR to amplify the *ORF50* promoter region up to 850nt upstream of the *ORF50* TSS. While PROX1 bound to the proximal regions of the *ORF50* promoter (up to 600nt upstream of the TSS) during the lytic cycle, (Fig.6f), it did not bind during latency (Supplementary Fig.5j). PROX1 binding to the *ORF50* promoter was also confirmed in KLECs (Fig.6f, lower panel).

To investigate the co-expression of PROX1 and viral lytic protein we stained consecutive sections of additional 36 KS biopsies (31 AIDS-KS and 5 AIDS-negative KS of which 26 from skin/lymph node lesions (LN) and 10 from gastro-intestinal (GI) lesions) with PROX1 and K8.1 antibodies, and observed that 92% of the skin/LN cases (24/26) and 50% (5/10) of the GI-KS were co-expressing PROX1 and K8.1 (Fig.6g). This suggests that PROX1 expression is associated with KSHV lytic protein expression also in KS tumours.

This data indicates that PROX1 enhances KSHV lytic gene expression, binds to the initiator of the KSHV lytic cycle, ORF50, and during the lytic cycle it binds to the *ORF50* promoter. PROX1 also enhances the auto-activation of *ORF50* promoter in an ORF50-dependent manner. In addition, in the majority of the analysed skin KS tumours PROX1 expression coincided with the expression of the late lytic marker K8.1, thus suggesting an important role for PROX1 in enhancing KSHV lytic gene expression in KS.

## DISCUSSION

KSHV infection of BECs or LECs results to different viral expression programs in the infected cells. While the virus is strictly latent in KBECs, it displays a unique, spontaneous lytic replication program in KLECs^5^. Multiple evidence supports a lymphatic rather than blood endothelial origin for KS-SC^3^. Our findings that SOX18, PROX1 and COUPTF2 are all expressed in KS-SC further support the lymphatic origin of KS-SC. We found that SOX18 and PROX1, both essential for maintaining the LEC identity, regulate KSHV life cycle through two different mechanisms in KLECs. COUPTF2 instead, although also expressed in KS, had no role in KSHV replication in KLECs.

SOX18 is listed among the 1482 genes differentially regulated in KS compared to normal skin^17^, but its role in the KSHV biology has been little explored. Here we show that SOX18 binds to the KSHV episome and increases their number. This is the first time that SOX18 is associated to a key function in viral life cycle. It has so far been linked to the transcriptional regulation of ECs and to the progression of solid cancers^34, 35^. SOX18 forms a dimer^20, 21^, able to bind to the viral and cellular genomes. This may lead to a stronger viral episome tethering to the host genome, thus improving the efficiency of KSHV genome retention. Supporting this hypothesis is the finding that SOX18 chemical inhibition of homo and heterodimerization reduced KSHV genomes. While the less characterized SM4 molecule will need further testing, the enantiopure R propranolol has already been proven safe^36^ and could in principle be repurposed to treat KS patients.

PROX1 has been studied in KSHV-induced EC reprogramming but little is known about its role in the KSHV life cycle. We found that PROX1 acts at the initiation of KSHV lytic cascade by enhancing transcription from the *ORF50* promoter in an ORF50-dependent manner. However, we do not exclude that PROX1 might also govern the expression of cellular regulator(s) of lytic reactivation and/or interact with other viral/cellular products and thereby enhance also other phases of the lytic cycle.

We further show that the late lytic protein K8.1 is expressed in the majority of the studied KS biopsies, suggesting that, similar to KLECs in culture, lytic protein expression is likely a common feature of KS tumours. This is supported by detection of the lytic proteins K5 and K15 in KS biopsies^37, 38^ and a recent RNA-seq study on African-KS lesions where KSHV lytic transcripts were detected in the tumours analysed^15^. It is not known whether KS-SCs, like the KLEC cultures, constitutively produce and release infectious virus.

KS is an atypical tumour: the SCs are diploid, depend on extracellular supply of growth factors for their survival, and viral persistence is necessary to support tumourigenesis^2, 3, 39, 40^. However, KSHV genome maintenance is inefficient and viral genomes are lost during cell division^41, 42^. Sporadic lytic reactivation and release of infectious virus is required *in vivo* to replenish the population of KSHV-infected SCs. Therefore, SOX18 and PROX1, as positive regulators of viral episome copies and productive lytic replication, are important not only for the KSHV life cycle but also for KS pathogenesis. Differently from the non-infected LECs, in KLECs PROX1, SOX18 and COUPTF2 expression are not interdependent. KSHV infection maintains high levels of these TFs and, in turn, PROX1 and SOX18 sustain viral infection. This suggests that the virus renders its cellular microenvironment optimal for its own persistence.

Overall, our findings revealed KLECs as a faithful *in vitro* model for KS research and answer some long-standing questions on the physiological control of KSHV spontaneous lytic gene expression pointing to endothelial TFs as important players in this process.

## METHODS

### Human subjects

Paraffin embedded KS sections were kindly provided by Dr. Justin Weir (Charing Cross Hospital, and the London Clinic, London). All KS cases analysed in the study were obtained from adult males. The study was covered by Riverside Research Ethics Committee (Study title: Kaposi’s Sarcoma Herpes Virus Infection And Immunity, REC reference: 04/Q0401/80); all patients gave written informed consent.

FFPE KS-negative skin biopsies were obtained from the archives of the Department of Pathology, Helsinki University Hospital, according to the Finnish laws and regulations by permission of the director of the health care unit. The tissue samples were de-identified and analysed anonymously.

### Cell culture

Primary human dermal lymphatic (C-12216) and blood (C-12211) endothelial cells (LECs and BECs) were purchased from Promocell. Each experiment was repeated using cells from at least two different donors.

Primary human LECs and BECs were grown in Lonza EBM-2 (00190860) supplemented with EGM^TM^-2 MV Microvascular Endothelial SingleQuots^TM^(CC-4147). HEK293FT (Thermofisher Scientific, R70007), U2OS (ATCC:HTB-96), HeLa (ATCC^®^ CCL-2^™^)and iSLK.219 were grown in DMEM, supplemented with 10% foetal calf serum, 1%L-glutamate, 1% pen/strep. iSLK.219 were also supplied with 10 ug/ml puromycin, 600μg/mL hygromycin B, 400μg/mL G418.

TREx BCBL1-Rta^43^ were grown in RPMI1640 supplemented with 20% foetal calf serum, 1%L-glutamate, 1% pen/strep.

### rKSHV.219 or WT KSHV production

rKSHV.219 was produced from iSLK.219 cells reactivated using 0.2 µg/ml doxycycline and 1.35mM NaB for 72h. WT KSHV was produced from TREx BCBL1-Rta^43^ reactivated with dox(1µg/ml) for 96h. Supernatant was then harvested, spun down (2000 rpm 5min) and sterile filtered using 45 µm pore-size filters. Subsequently the supernatant was ultracentrifuged at 22000 rpm for 2h. The concentrated virus was then aliquoted and stored at -80°C.

Virus titers were determined by infecting U2OS cells with serial dilutions of the concentrated virus preparation and assessing the amount of GFP+ or LANA+ cells 24h post-infection by automated high-content microscopy (see below in the virus titration section).

### KSHV infection of endothelial cells

Early passage LECs or BECs (less than 6) were grown to semi-confluency and infected at MOI of 1.5 and 3, respectively (to achieve the same number of virus positive cells) by spinoculation (450g, 30 min RT) in the presence of 8µg/ml of polybrene. Media was replaced every third day. At 7-9 d.p.i. cells were split at 1:2 or 1:3 ratio and the experiments started after additional 2-4 days when cells reached again confluency and organized spindling and RFP expression (as a marker of lytic expression) were observed.

### Plasmids and lentivirus constructs

The following plasmids were used in this study: pAMC Prox1 WT, pAMC Prox1 MUT, pAMC (described in ^31^); pcDNA ORF50 (described in ^44^); pcDNA LANA; pcDNA 3.1 (Thermo Fisher); pGL2-ORF50 (described in ^44^); pGL2-basic (Promega); pGL3-7XTR (provided by TF Schulz, Hannover Medical School) and pGL3-basic (Promega).

PROX1 WT and PROX1 MUT expressing lentiviruses were described previously^20^. The control pFuW-myc-NLS-mCherry was generated from the pFuW-myc-BirA-NLS-mCherry (a kind gift by R.Kivela, University of Helsinki). SOX18 gene sequence was codon optimized and synthesized by GeneArt (Thermo Fisher). The codon optimized SOX18 was cloned in the pFuW-myc by Gibson Assembly using as an initial backbone pFuW-myc-NLS-mCherry. Backbone and insert were assembled using the NEB HiFi DNA assembly (New England BioLabs, E2611).

Inserts were verified by Sanger sequencing and the integrity of the backbone by restriction analysis.

### Lentivirus production and transduction

5×10^6^ HEK293FT cells were plated in a T75 flask. Next day cells were transfected with 3^rd^ generation lentivirus packaging plasmids (2.83 µg pLP1, 1.33 µg pLP2, 1.84 µg pLP/VSVg) and 5 µg of expression plasmid of interest using 20 µl/flask of Lipofectamine 2000. Next day, media was changed and the virus was collected 48h later. The lentivirus was then concentrated using PEG-IT (System Biosciences) according to the manufacturer instructions.

iSLK.219 cells were plated for transduction at a density of 2.5×10^5^ cells/ml. On the next day transduction was done by spinoculation (450g, 30 min, RT) in the presence of 8µg/ml of polybrene. On the next day, cells were supplied with fresh media.

### Plasmids and siRNAs transfections

DNA transient transfections were performed using Lipofectamine 2000 (Invitrogen) for lentivirus preparation (see above) and FuGENE XD (Promega) for luciferase reporter assays at a ratio 1µg DNA:4 µl FuGENE XD.

Transient transfection of siRNA in a semi-confluent culture of rKSHVLEC was done using 3 µl of Lipofectamine RNAiMAX (Invitrogen) and 150pmol siRNA in a 6 well plate and following manufacturer instructions. Next day cells were supplied with fresh media. The following siRNA - were used: Stealth RNAi™ Prox1-1 and 2 (HSS 108596; HSS 108597) and Stealth RNAi™ Negative Control (HSS 12935200) from Invitrogen; ON-TARGETplus SOX18 siRNA (L-019035-00); ON-TARGETplus COUPTF2siRNA (L-003422-00) and ON-TARGETplus Non-targeting pool siRNA (D-001810-10) from Dharmacon.

### Virus titration

One day prior titration 8000 U2OS/well were plated on viewPlate-96black (6005182, Perkin Elmer). Cells were infected in triplicate by spinoculation (450g, 30 min, RT) in the presence of 8µg/ml of polybrene with serial dilution of pre-cleared (by centrifugation 8-10 min at 1000g) supernatant from cells treated as indicated. Next day cells were fixed in 4%PFA, stained with GFP (a kind gift from J. Mercer; UCL, UK) or LANA (Rat monoclonal HHV-8 (LN-35) ab4103; Abcam) and Hoechst 44432 (1 µg/ml).

Plates were imaged using Thermo-Scientific Cell Insight High Content Screening (HCS) system; images from 9 fields/well were taken.

### Inhibitor treatments

PAA (Cat#284270); R-Propranolol (Cat#P0689), S-Propranolol (Cat#P8688) and R-S Propanolol (Cat#P0884) were obtained from Sigma-Aldrich; Sm4 was kindly provided by M. Francois^20^.

KLECs or iSLK.219 cells were treated for six and two days, respectively with the indicated inhibitor at the concentrations stated in the Figure panels prior to the experiments.

During the 6-day drug treatment, the media was changed with fresh media-containing drug every 2 days.

### Luciferase Reporter assay

2.5×10^5^ HEK293FT/ml were plated in 24well plates (500ml/well). Next day each well was transfected using FuGENE XD (Promega) with 0.1 µg of reporter plasmid (or the corresponding vector controls), 0.25 µg of RTA (or the corresponding vector control), 0.25 µg of the plasmid of interest (PROX1 WT or MUT). Alternatively, HeLa cells were transfected with 0.1 µg of reporter plasmid (or the corresponding vector controls), 0.2 µg of LANA (or the corresponding vector control), 0.25, 0.5 or 1 µg of SOX18-expressing plasmid or the mCherry vector control. 32-36h post-transfection cells were lysed in 1Xpassive lysis buffer (Promega). Experiments were done at least two times in duplicates, bars represent the average and the error bars the SD across the different experiments.

### Quantification of virus genome copy number

Equal number of cells (10^6^) were harvested and cellular and viral DNA was isolated with NucleoSpin Tissue Kit (Macherey-Nagel, Cat#740988). The extracted DNA was then analysed by qPCR using primers for genomic actin (AGAAAATCTGGCACCACACC; AACGGCAGAAGAGAGAACCA) and K8.1 (AAAGCGTCCAGGCCACCACAGA; GGCAGAAAATGGCACACGGTTAC).

### Co-immunoprecipitation (Co-IP)

iSLK.219 cells were transduced with Myc-tagged PROX1 (WT or MUT) lentiviral vectors or an empty pSIN control vector. 24h later cells were reactivated with 0.2 µg/ml doxycycline and one day later cells were lysed in IP buffer (150 mM NaCl, 25 mM tris HCl pH 7,6, 1 mM EDTA, 1% glycerol, 0.2% NP-40).

Pre-cleared lysates were incubated with either mouse monoclonal Myc-tag (1:100) or normal mouse IgG (1:100) antibodies O/N at 4°C with gentle rotation and, subsequently, 30 µl of pre-washed Protein G sepharose beads (ab 193259, Abcam) were added and incubated for 2h at 4°C with gentle rotation. Beads were then washed 3 times in ice-cold IP buffer and the protein complexes analysed by western blot. Experiments were done at least two independent times, representative, cropped blots are shown, uncropped blots are shown in Supplementary Fig.6.

### Chromatin-immunoprecipitation (ChIP)

For each ChIP one 10 cm dish of iSLK.219 cells transduced with the vector of interest were used. The experiments were performed using the Simple ChIP enzymatic chromatin IP kit (Cell Signaling Technology, cat# 9003S). For chromatin IP the following antibodies were used: rabbit polyclonal anti-PROX1 (Cat#11067-2-AP, Proteintech Group), mouse monoclonal anti-Myc (Cat#2276, Cell Signaling Technology), normal mouse or rabbit IgG (sc-2025 and sc-3888, Santa cruz biotechnology), mouse monoclonal anti-HA.11(16B12, BD Pharmingen).

The experiments were done at least two independent times, Graphs show the average across the different experiments ±SEM (Fig.5) or SD (Fig.6).

The isolated DNA was amplified with the following primers:

ORF50 -620-850 (F;R) gtggtagagccagcagacgttc; tgtagcgccatctctgccc

ORF50 -320-610 (F;R) gggtgatttcttctaccacggtcat; ccgagcgtattctcagaggtct

ORF50 0-320 (F;R) tggcattttgttgcgcgcatgatc; ccgagcgtattctcagaggtc

OriA1 (F;R) ctccccggcaacaacctg; gggggttatatgcgcgtgc

OriA2 (F;R) caagcacgcgcatataaccc; gggatatgcttccgcctcat

OriA3 (F;R) caccgtgttagtgtcaccca; caccgtgttagtgtcaccca

Orilyt1 (F;R) attcaaagggggcacagagg; atgctgggacagaatagccg

Orilyt 2 (F;R) tctgtcccagcataggctc; cctgtgcccaaatctgtcct

Orilyt 3 (F;R) cacgcgggttgtttgaaagt; ccacttgggtgcacagagat

TR1 (F;R) cataaatattccggatacaaggctcg; gactcctcgcacagtagagagag

TR2 (F;R) actgacaaacaaaatgcacataacaag; actgacaaacaaaatgcacataacaag

TR3 (F;R) gaacatcagggatgggtctatgatc; gataaccctcacctaccatggaaat

TR4 (F;R) gataaccctcacctaccatggaaat; agagctacgagtgtcataaatacaaga

### Immunohistochemistry

After deparaffinization in xylene and rehydration, for antigen retrieval, sections were treated in a PreTreatment module (Lab Vision Corp., Fremont, CA, USA) in Tris-HCl (pH 8.5) and in Tris-EDTA (pH 9.0) buffer for 20 minutes at 98°C. Sections were stained in an Autostainer 480 (Lab Vision). Tissues were incubated with the indicated antibodies (mouse anti Sox18, 1:100; mouse anti COUPTF2, 1:150; mouse anti K8.1, 1:250) overnight at room temperature. For detection, ImmPRESS HRP Polymer Detection Kit, Peroxidase, (Vector Laboratories, Burlingame, CA, USA) was used.

LANA staining was done O/N at 4 °C in humidified chambers using rabbit polyclonal antibody (a kind gift from B. Chandran, University of South Florida) after washing 3 times the slides in PBS-T, sections were stained with goat anti-rabbit Alexa594 conjugated antibody for 1 hour at RT, nuclei were counterstained with Hoechst 33342 (1ug/ml).

Samples were imaged in a Panoramic 250 viewer (Genome Biology Unit, Research Programs Unit, University of Helsinki), representative snapshots are shown.

### RNA-sequencing (RNA-seq)

RNA from iSLK.219 cells and KLECs treated as indicated and from 3 independent experiments, was extracted using NucleoSpin RNA extraction kit (Macherey Nagel). Ribosomal RNA (rRNA) was depleted using ribo-zero rRNA removal kit (Illumina) and the quality of the samples was monitored with Bioanalyzer RNA Kit (Agilent). Libraries were prepared using NEB Next Ultra Directional RNA library Prep Kit for Illumina (New England BioLabs) and sequencing was done with NextSeq High Output 1X75bp (75SE).

FASTQ data was aligned to the annotated human reference genome hg38 and using STAR ^45^with the build in gene quantification function. Differential gene expression (DGE) of the quantified genes was performed using DeSeq2 ^46^. Significantly DGE (FDR < 0.1 and log2FoldChange >= +/- 1) were subjected to Kyoto Encyclopedia of Genes and Genomes (KEGG) pathway analysis to identify regulated pathways (FDR < 0.1). Concerning viral gene expression differences FASTQ data was aligned to the rKSHV.219 reference sequence which is identical to BAC16 (Genbank Acc.: GQ994935) using STAR. We inserted the panPromoter-RFP, GFP and PuroR cassette according to the original publication into the reference for analysis ^19^, since it was not part of the deposited Genbank sequence. Read counting of viral ORFs was performed using FeatureCounts ^47^. KSHV specific counts were normalized using the estimated size factors generate for the human dataset to correct for sequencing depth-based bias. DGE for KSHV genes was performed using DeSeq2.

To detect KSHV- encoded circular RNAs, FASTQ files were aligned using STAR to the human (hg19) + KSHV hybrid genome file. CIRCexplorer2 was used to detect back-spliced junctions in the KSHV genome. All back-spliced junctions were mapped in 200 nucleotide-sized bins and plotted.

RNA-seq raw data have been deposited in the European nucleotide Archive (ENA) under the ID code: ena-STUDY-UNIVERSITY OF HELSINKI-13-02-2019-22:25:30:309-452 and the accession number: PRJEB31253.

Data available at the following URL: http://www.ebi.ac.uk/ena/data/view/PRJEB31253.

### RT-qPCR

Total RNA was extracted from cells using NucleoSpin RNA extraction kit (Macherey Nagel) and reverse transcribed as described in^11^. Transcripts were measured using 2XSYBR reaction mix (Fermentas) and unlabeled primers (F;R) listed below:

PROX1 (F;R) TGTTCACCAGCACACCCGCC; TCCTTCCTGCATTGCACTTCCCG;

SOX18 (F;R) CTTCATGGTGTGGGCAAAG; GCGTTCAGCTCCTTCCAC;

COUPTF2(F;R) GCAAGTGGAGAAGCTCAAGG; TCCACATGGGCTACATCAGA;

Actin(F;R) TCACCCACACTGTGCCATCTACGA; TCACCCACACTGTGCCATCTACGA;

Orf73(F;R) ACTGAACACACGGACAACGG; CAGGTTCTCCCATCGACGA;

Orf50 (F;R) CACAAAAATGGCGCAAGATGA; TGGTAGAGTTGGGCCTTCAGTT;

Orf 57 (F;R) TGGACATTATGAAGGGCATCCTA; CGGGTTCGGACAATTGCT;

Orf45 (F;R) CCTCGTCGTCTGAAGGTGA; GGGATGGGTTAGTCAGGATG;

K8.1 (F;R) AAAGCGTCCAGGCCACCACAGA; GGCAGAAAATGGCACACGGTTAC

Each sample was measured in triplicate and normalized to the *Actin* levels. Experiments were done at least two independent times, the graphs show the average and error bars the SD across the experiments.

### Western Blotting and antibodies

Cells were lysed in RIPA buffer (150 mM NaCl, 1% Igepal CA630, 0.5% Na deoxycholate, 0.1% SDS, 50 mM Tris-HCl-PH 8.0), supplemented with protease and phosphatase (Pierce™ 88666, 88667). After pre-clearing by centrifugation, lysates were mixed with 5X laemli sample buffer and run for 40 min at 55mA on Criterion TGX precast gels (Bio-Rad).

Transfer on nitrocellulose membrane was done using trans-blot Turbo Transfer system (Bio-Rad). Blocking was performed in 5% non-fat dry milk in TBS-T (0.1% Tween) for 1h RT. Membranes were incubated in primary antibody O/N 4°C (either 5% BSA or non-fat dry milk in TBS-T, according to the primary antibody manufacturer’s recommendation). The following primary antibodies were used: Mouse monoclonal anti- actin (Santa Cruz Biotechnology, sc-8432); Mouse monoclonal anti-gamma-tubulin (Sigma-Aldrich, T6557); Rabbit monoclonal anti-PROX1 (Cell Signaling Technology D2J6J); Mouse monoclonal anti SOX18 (D-8) (Santa Cruz Biotechnology, sc-166025); Mouse monoclonal anti COUP-TFII (Perseus Proteomics, PP-H7147-00); Rat monoclonal HHV-8 (LN-35) (Abcam, ab4103) Mouse monoclonal anti KSHV ORF57 (Santa Cruz Biotechnology, sc-135746); Mouse monoclonal anti KSHV ORF45 (Santa Cruz Biotechnology, sc-53883); Mouse monoclonal anti KSHV K8.1 (Santa Cruz Biotechnology sc-65446); Rabbit monoclonal anti GFP (Cell Signaling Technology, 2956); Rabbit polyclonal anti ORF50 (Kind gift from C. Arias).

After three rounds of washing in TBS-T, membranes were incubated in the appropriate HRP-conjugated secondary antibody (from Cell Signaling Technology) diluted in blocking solution for 4h RT. Luminescent signal was revealed with Wester-Bright Sirius detection Kit (Advansta). Experiments were done at least two independent times, representative experiments are shown. Cropped immunoblots are presented in each figure, the corresponding uncropped membranes are presented in Supplementary Fig.6.

The Fiji software (https://imagej.net/Fiji) was used to quantify the intensity of the bands. For each sample, band intensities relative to the corresponding loading control is shown.

### Immunofluorescence

2×10^4/ml primary BEC and LEC infected or not with rKSHV.219 were plated on coverslips for 4 days. 10^3 iSLK.219 were plated on viewPlate-96 black (6005182, Perkin Elmer) transduced as indicated after 24h and, one day later reactivated with Doxycycline for 24h.

Coverslips and plates were fixed in 4%PFA 20 min RT, permeabilized (0.3% tritonX in PBS) and stained with Hoechst 33342 (1 µg/ml) 10 min RT. Blocking, washings and antibody incubations were done in 0.5%BSA in PBS. Coverslips were incubated in primary and, after washing, in secondary antibody in humidified chambers for 2h RT and then mounted in Mowiol. Experiments were done at least two independent times. The following antibodies were used: Goat polyclonal anti-PROX1 R&D Systems, AF2727); Mouse monoclonal anti SOX18 (D-8) (Santa Cruz Biotechnology, sc-166025); Mouse monoclonal anti COUP-TFII(Perseus Proteomics, PP-H7147-00); Rabbit polyclonal anti LANA (kind gift from B. Chandran); Mouse monoclonal anti KSHV ORF57 (Santa Cruz Biotechnology, sc-135746); Mouse monoclonal anti KSHV K8.1 (Santa Cruz Biotechnology sc-65446); Rabbit polyclonal anti ORF50 (a kind gift from C. Arias, University of California, Santa Barbara?). All secondary antibodies were from Thermo Fischer Scientific: Goat anti-Rabbit IgG (H+L) (A-11034); Goat anti-Rabbit IgG (H+L) Highly Cross-Adsorbed (A32733); Goat anti-Mouse IgG (H+L) Cross-Adsorbed (A32728); Goat anti-rat IgG (H+L) Cross-Adsorbed Alexa Fluor 647 (A-21247).

Coverslips were imaged in Zeiss LSM 780 confocal microscope. Plates were imaged using Thermo Scientific Cell Insight High Content Screening (HCS) system.

### Analysis of cell proliferation

Cells were maintained for 1 h in media containing 10 µM 5-ethynyl-2’-deoxyuridine (EdU) and fixed in 4% paraformaldehyde in PBS. The EdU-incorporated in the cell DNA was coupled to Alexa Fluor 647 as per instructions provided in the Click-iT EdU Alexa Fluor 647 Imaging Kit (Thermo Fischer Scientific). Images were acquired using a CellInsight High Content microscope (Thermo Fischer Scientific).

The portion of EdU containing cells was quantified using the Cell Profiler software.

### Quantification of virus titers, lytic cells and fluorescent signal intensity

The number of positive cells were quantified over the total amount of cells (number of nuclei) using Cell Profiler (http://cellprofiler.org). The graph shows the average of virus titers or positive cells per each condition, error bars indicate SD across at least two experiments. The virus titers were calculated as IU/ml. Average signal intensity was calculated with appropriate cell profiler pipeline, the graph shows the average intensity per condition and the error bars indicates the SD across n>100 cells/condition/staining.

Co-localization was assessed by Pearson’s co-localization coefficient calculated using Coloc2 plug-in in Fiji software package (https://imagej.net/Fiji).

### Statistical analysis

Experimental data were plotted and analysed with Prism GraphPad versions 7 and 8. Unless differently stated in the legend, the graphs show the single values of each biological replicate (for n<20), the mean and the error bars indicate the SD across the biological replicates.

Ordinary one-way Anova followed by Dunnett’s correction for multiple comparisons or or by two-tailed paired t-test were performed to assess the statistical significance of the differences between the different treatments and the control sample. The significance is shown in the Figures as follows: *: p<0.033; **: p<0.02, ***: p<0.001. Exact P values are indicated in the appropriate graphs or in the Supplementary Table3.

### Data availability

The data that support the findings of this study are available from the corresponding author upon reasonable request.

RNA-seq raw data have been deposited in the European nucleotide Archive (ENA) under the ID code: ena-STUDY-UNIVERSITY OF HELSINKI-13-02-2019-22:25:30:309-452 and the accession number: PRJEB31253.

Data available at the following URL: http://www.ebi.ac.uk/ena/data/view/PRJEB31253.

## SUPPLEMENTARY MATERIAL

Supplementary Figures 1-6.

Supplementary Tables 1-3.

## ACKNOWLEDGEMENTS

We are grateful to Paul Farrell for critical comments on the manuscript and to Takanobu Takawa for the technical support in the detection of Kcirc RNAs. We thank the Biomedicum Imaging Unit (BIU) and Genome Biology Unit (GBU) at the University of Helsinki for support in imaging as well as the Functional Genomic Unit (FuGU, University of Helsinki) for technical support during the RNA-seq. Dr. Justin Weir (Imperial College Healthcare NHS Trust) is acknowledged for kindly providing the KS biopsies. We are also extremely grateful to Nadezhda Zinovkina and Pia Foxell (University of Helsinki) for the valuable technical help. Bala Chandran (University of South Florida) and Carolina Arias (University of California Santa Barbara) are thanked for antibodies to LANA and ORF50, respectively.

The study was supported by the Academy of Finland Centre of Excellence grant (Translational Cancer Biology; PMO) and the Academy of Finland postdoctoral researcher grant (SG), Finnish Cancer Foundations (P.M.O) and Sigrid Juselius Foundation (P.M.O.).

## AUTHOR CONTRIBUTIONS

Conceptualization: SG, EE, PMO; Methodology: SG, EE; Validation: SG, KT; Formal Analysis: SG, EE, KT, TG; Investigation: SG, KT, VN, TG, REK, RD; Resources: SPK, MB, CH, MF; Data Curation: SG, TG, AG; Writing- Original Draft: SG, PMO; Writing- Review & editing: SG, EE, AG, JMZ, MB, MF, PMO; Visualization: SG, PMO; Supervision: SG, PMO; Project Administration: PMO; Funding Acquisition: SG, PMO.

