## Supplementary figures 1-6 for "Oncogenic herpesvirus engages the endothelial transcription factors SOX18 and PROX1 to increase viral genome copies and virus production"

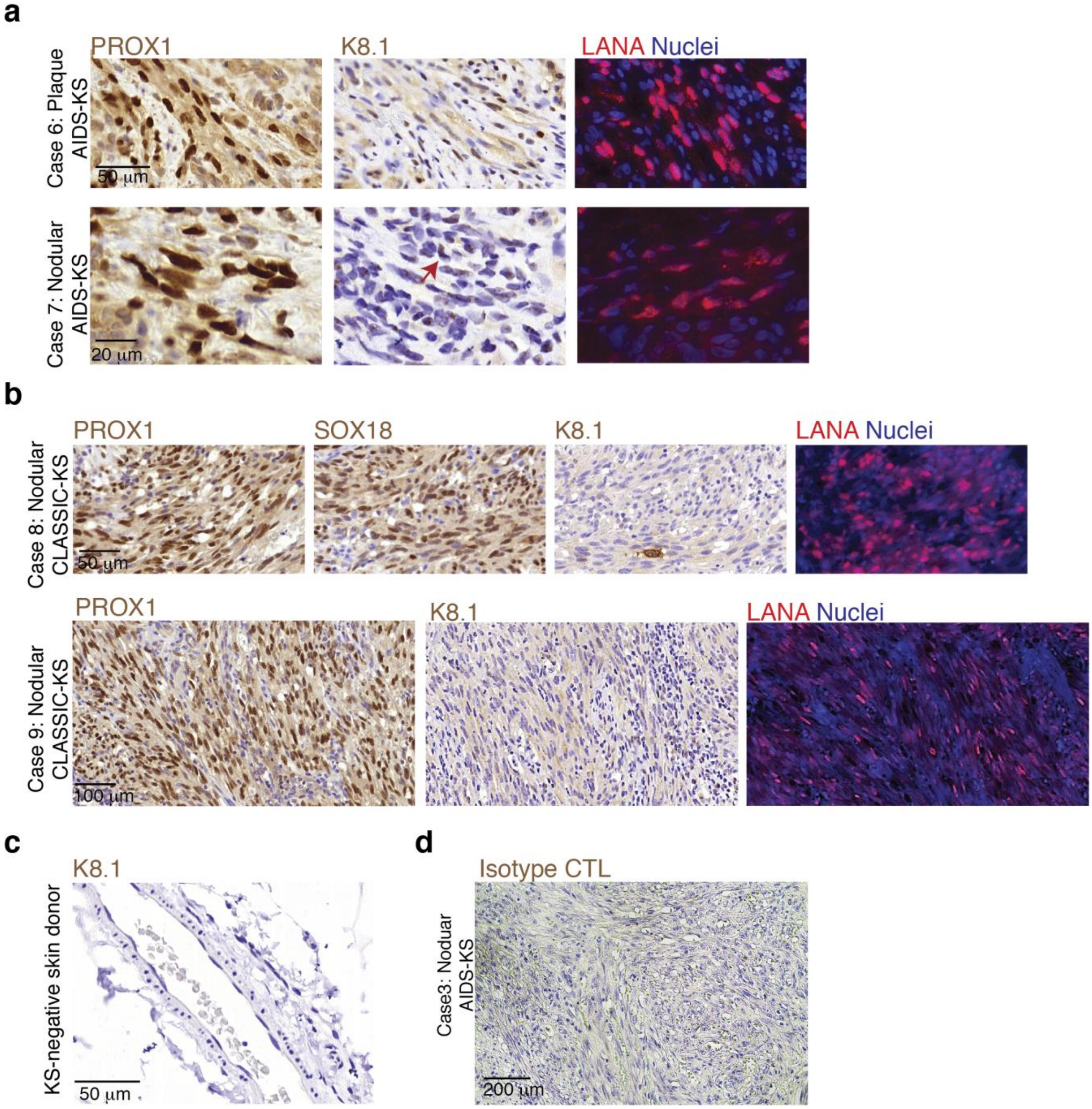


**Supplementary Fig. 1. (a-b)** Representative images of consecutive sections from (a) 2 AIDS-KS patients and (b) 2 classic KS patients stained with the indicated antibodies. The arrow indicates a cell positive for K8.1 staining. **(c)** KS negative skin donor stained with the K8.1 antibody. **(d)** Representative image of an AIDS-KS biopsy stained with an isotype control (CTL) antibody for K8.1.

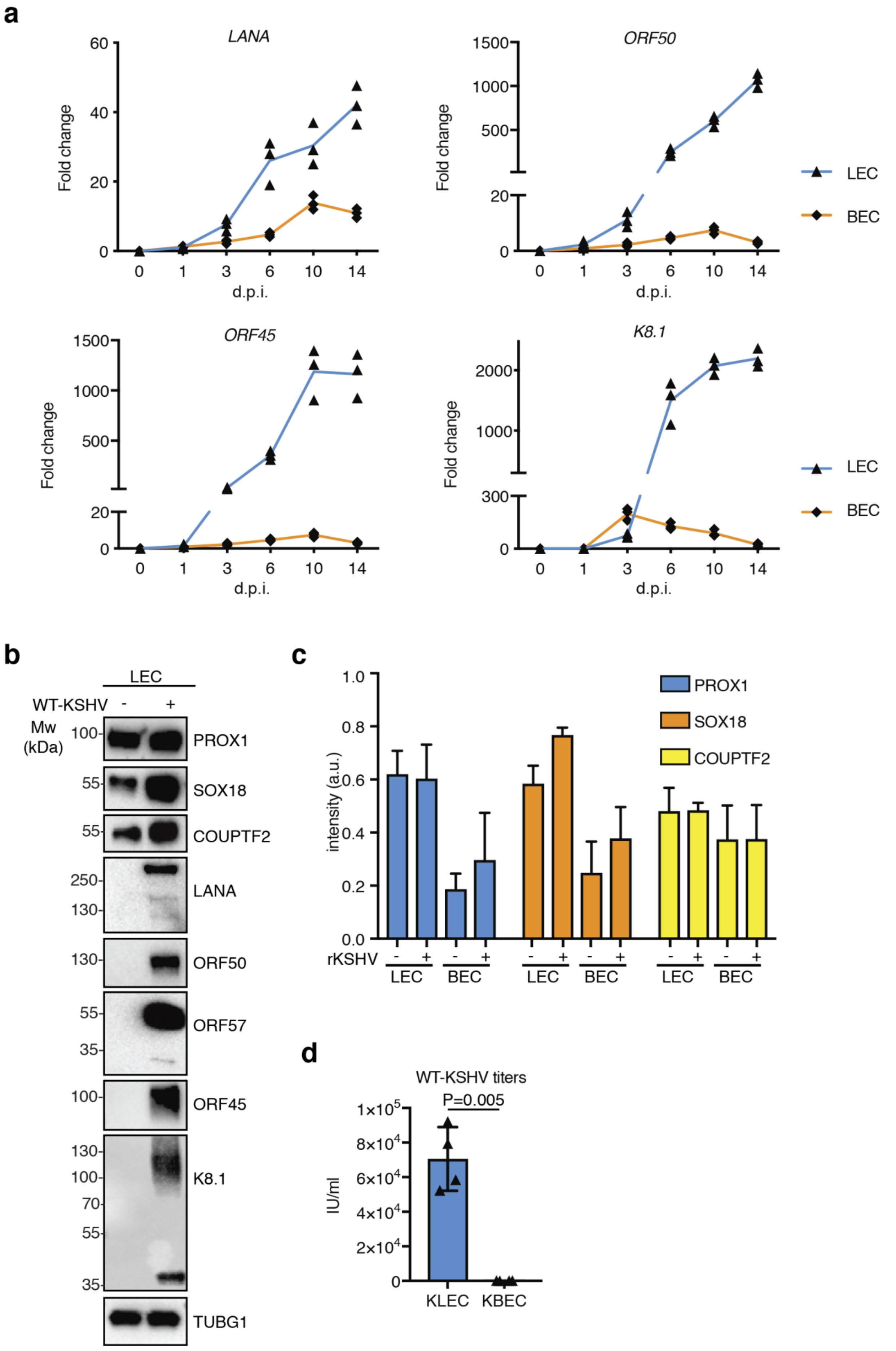

a
LANA
ORF50
ORF45
K8.1
Fold change
d.p.i.
LEC
BEC
b
LEC
WT-KSHV - +
Mw (kDa)
PROX1
SOX18
COUPTF2
LANA
ORF50
ORF57
ORF45
K8.1
TUBG1
c
intensity (a.u.)
PROX1
SOX18
COUPTF2
rKSHV
LEC
BEC
d
WT-KSHV titers
P=0.005
IU/ml
KLEC
KBEC

**Supplementary Fig. 2. (a)** KLECs and KBEC were infected with rKSHV.219 and cells were analysed at the indicated timepoints by RTqPCR for the indicated viral targets (d.p.i.: days post-infection). **(b)** LEC were infected with BCBL1-TreX-derived KSHV left uninfected; 14 d.p.i. cells were harvested and analysed by immunoblot for the expression of the indicated proteins; gamma-tubulin (TUBG1) as loading control. Representative, cropped immunoblots are shown, uncropped blots are shown in Supplementary Fig.6. The experiment was done two independent times. **(c)** quantification of the mean fluorescent intensity for the IF shown in Figure 2d. Bars represent mean ±SD of n>100 cells/condition/staining. a.u.: arbitrary units. **(e)** LEC and BEC were treated as in (b) and the infectious virus released in the supernatant was quantified. Single values from n=4 biological replicates are shown. Bars represent mean ± SD. P value was calculated using two-tailed paired t-test.

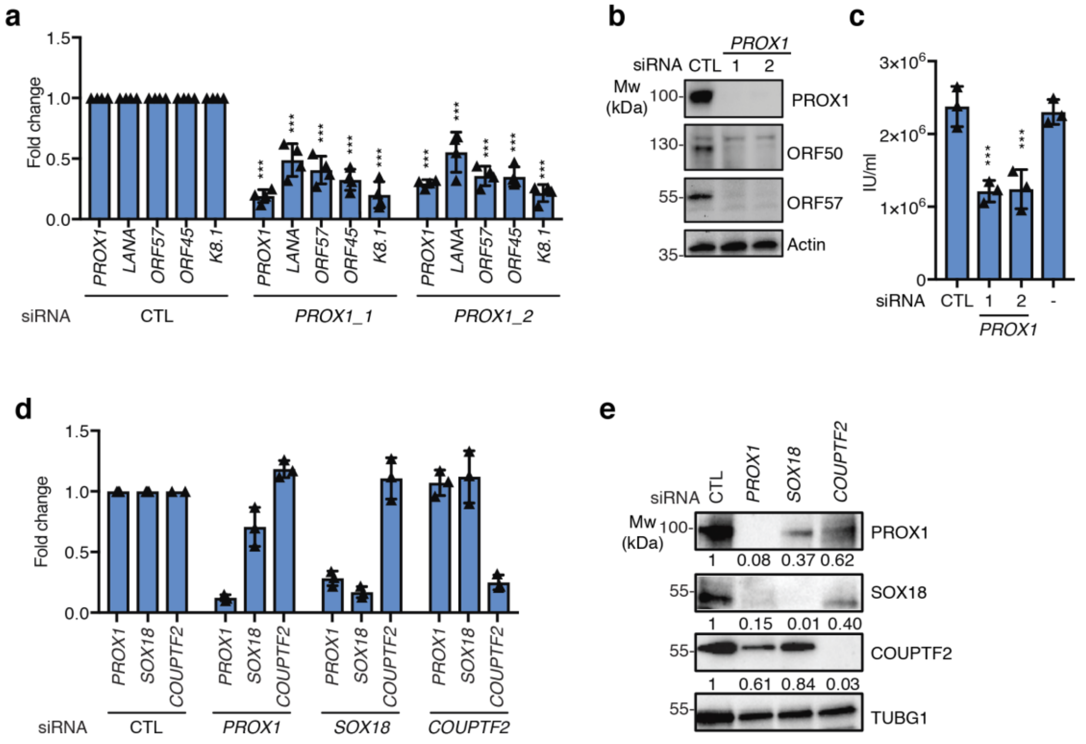


**Supplementary Fig. 3. (a-c)** KLEC were treated with control siRNA (CTL) or two different PROX1 targeting siRNAs for 72h. Cells were analysed (a) by RTqPCR for the indicated viral and cellular targets; single values from n=3 independent experiments are shown. Bars represent mean ± SD. (b) by immunoblot for the expression of the indicated proteins using actin as a loading control; the experiment was repeated two independent times. Representative, cropped immunoblots are shown, uncropped blots are shown in Supplementary Fig.6. The experiment was done two independent times. (c) the cell free supernatant was titrated for the presence of infectious virus. Single values from n=3 independent replicates are shown. Bars represent mean ± SD **(d)** The silencing efficiency of the experiment shown in Fig.3a was assessed by RTqPCR. Single values from n=3 independent replicates are shown. **(e)** LEC were treated with the indicated siRNAs for 72h. Cell lysates were analysed by immunoblot for the indicated targets, TUBG1 as a loading control. Representative, cropped immunoblots are shown, uncropped blots are shown in

Supplementary Fig.6. The experiment was done two independent times. Numbers indicate the quantified intensity of each band relative to the appropriate loading control.

P values in panels (a) and (c) were calculated using ordinary one way-anova followed by Dunn´s correction for multiple comparisons. *: $p<0.033$; **: $p<0.02$, ***: $p<0.001$. Exact p values are shown in Supplementary Table3.

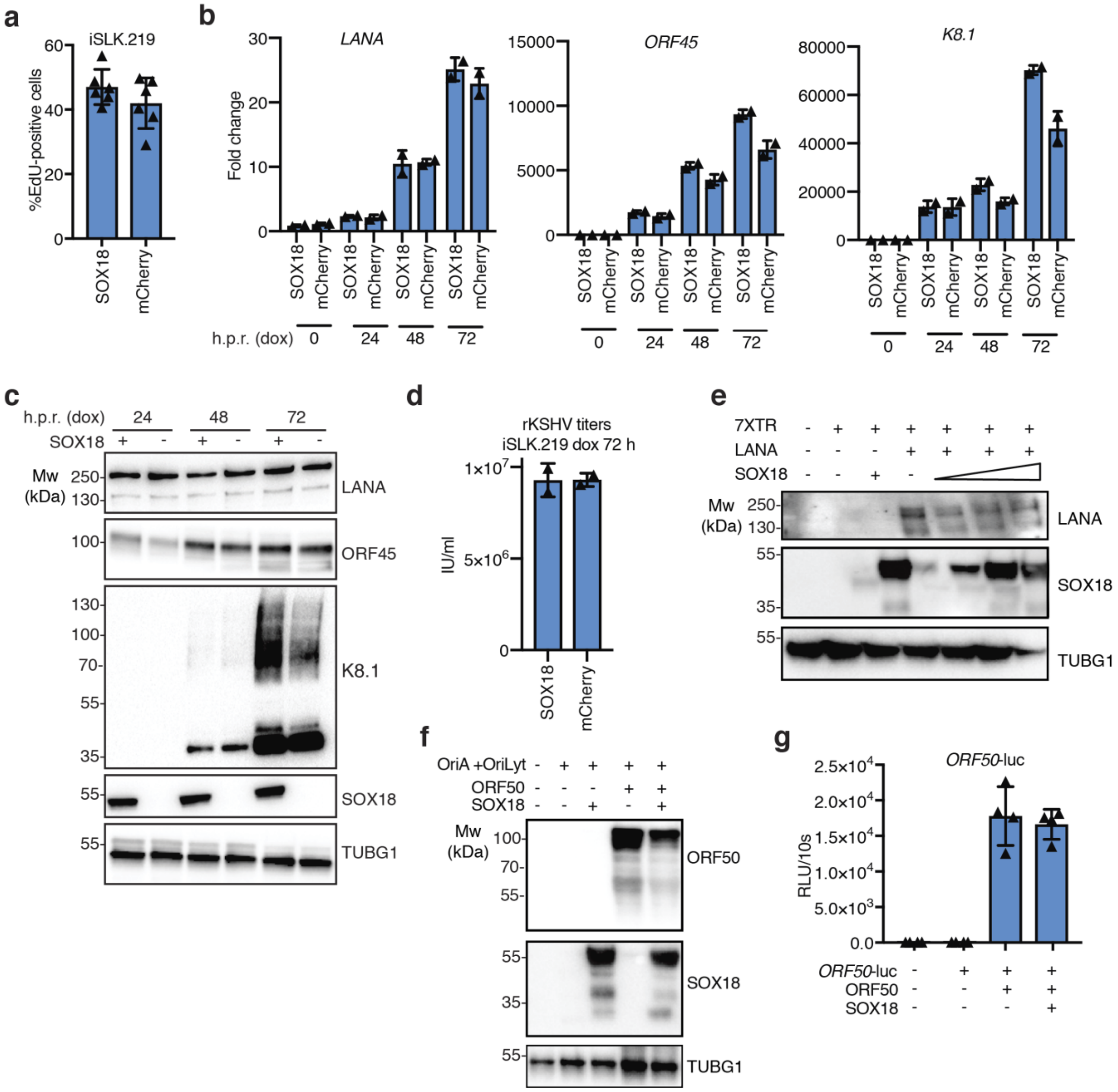


**Supplementary Fig. 4**. **(a)** iSLK.219 were transduced with SOX18- or mCherry-expressing lentiviruses and next day treated with EdU for 1h and then fixed. Percentage of EdU positive cells is shown for n=6 biological replicates, bars represent mean ± SD. **(b-d)** iSLK.219 were transduced as in (a) and reactivated with dox (h.p.r.: hours post reactivation). Cells were harvested at the indicated timepoints and analyzed (b) by RTqPCR, single values from n=2 independent experiments are shown. Bars represent mean ± SD (c) by immunoblot for the indicated viral proteins, gamma-tubulin (TUBG1) as a loading control. **(d)** Titration for released infectious virus from iSLK.219 transduced and treated as indicated. Single value from n=2 independent experiments are shown. Bars represent mean ± SD **(e,f)** Immunoblot analysis of the samples used for the luciferase assays in Figure 5b,c gamma-tubulin (TUBG1) as a loading control. Representative, cropped immunoblots are shown, uncropped blots are shown in Supplementary Fig.6. **(g)** Luciferase reporter assay in HEK293FT transduced as indicated. Single values for n=4 biological replicates are shown, bars represent mean ± SD.

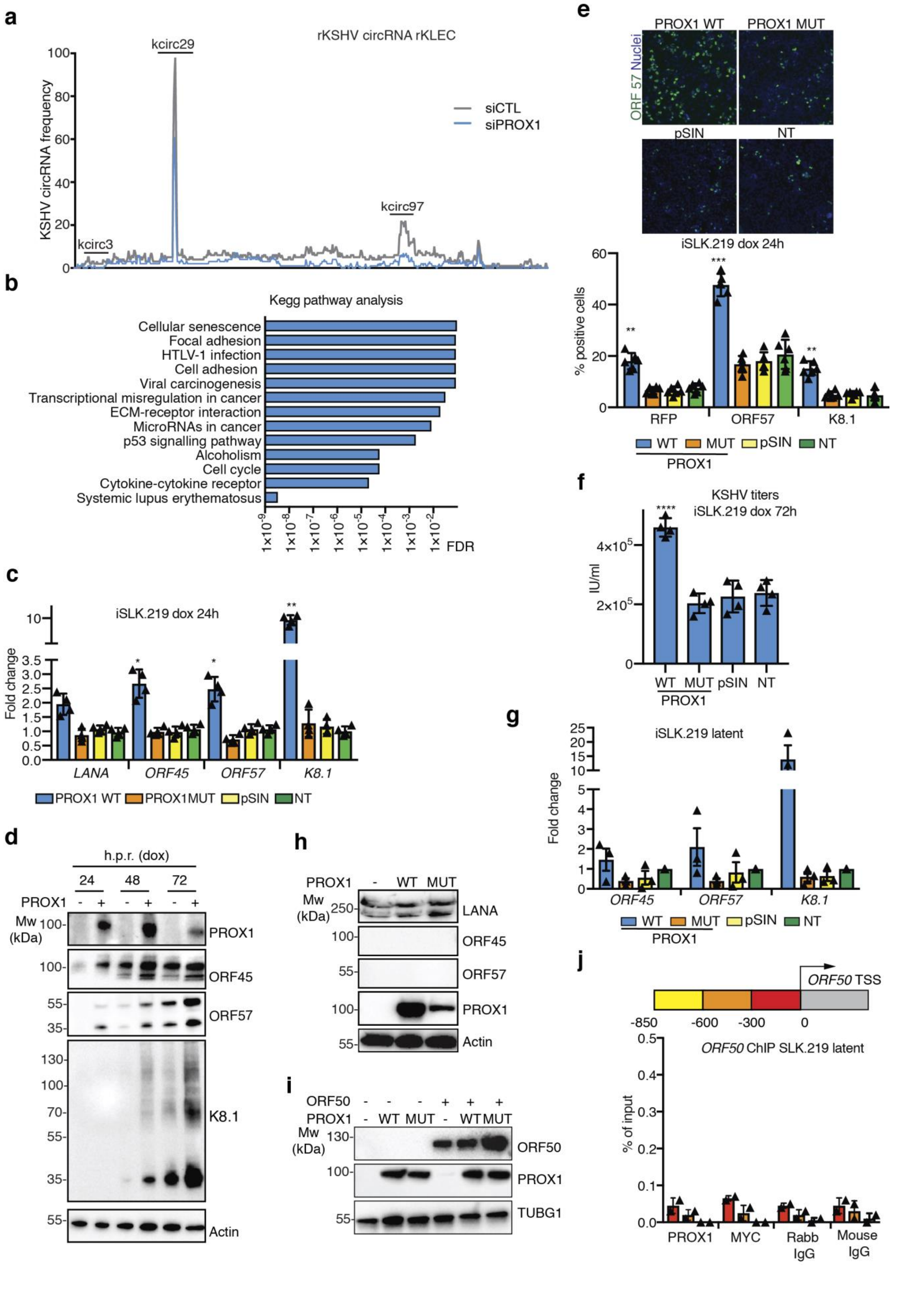
a
rKSHV circRNA rKLEC
KSHV circRNA frequency
kcirc29
kcirc3
kcirc97
siCTL
siPROX1
b
Kegg pathway analysis
Cellular senescence
Focal adhesion
HTLV-1 infection
Cell adhesion
Viral carcinogenesis
Transcriptional misregulation in cancer
ECM-receptor interaction
MicroRNAs in cancer
p53 signalling pathway
Alcoholism
Cell cycle
Cytokine-cytokine receptor
Systemic lupus erythematosus
FDR
c
iSLK.219 dox 24h
Fold change
LANA
ORF45
ORF57
K8.1
PROX1 WT
PROX1MUT
pSIN
NT
d
h.p.r. (dox)
PROX1
Mw (kDa)
PROX1
ORF45
ORF57
K8.1
Actin
e
PROX1 WT
PROX1 MUT
ORF 57 Nuclei
pSIN
NT
iSLK.219 dox 24h
% positive cells
RFP
ORF57
K8.1
WT
MUT
pSIN
NT
PROX1
f
KSHV titers
iSLK.219 dox 72h
IU/ml
g
iSLK.219 latent
Fold change
ORF45
ORF57
K8.1
h
PROX1
LANA
ORF45
ORF57
PROX1
Actin
i
ORF50
PROX1
ORF50
PROX1
TUBG1
j
ORF50 TSS
ORF50 ChIP SLK.219 latent
% of input
PROX1
MYC
Rabb IgG
Mouse IgG

**Supplementary Fig.5.** **(a)** frequency of KSHV-encoded circ-RNA assessed by RNA-seq in KLEC treated with control or PROX1 targeting siRNAs. (**b)** Significantly enriched pathways of DEG (differentially expressed genes) in rKLEC treated with siPROX vs siCTL and subjected to RNA-seq, identified by KEGG analysis pathway. FDR: false discovery rate. **(c-f)** iSLK.219 were transduced with lentiviral vectors expressing the indicated proteins or non-treated (NT) and one day later treated with dox and harvested at the indicated timepoints. (c) Cells were analyzed by RTqPCR for the indicated viral targets 24h after reactivation. Single values from n=3 independent experiments are shown. Bars represent mean ± SD. (d) Immunoblot for the indicated viral proteins, actin as a loading control. h.p.r.= hours post reactivation. Representative, cropped immunoblots are shown, uncropped blots are shown in Supplementary Fig.6. The experiment was done two independent times. (e) high content image analysis for the indicated markers 24h post-reactivation. Upper panels: representative images of ORF57 -positive cells in each condition, nuclei were counterstained with Hoechst 33342, Lower panel: average % of positive cells expressing indicated proteins/condition. Single values from n=6 biological replicates are shown. Bars represent mean ± SD. (f) Titration of released infectious virus from iSLK.219 transduced as indicated and reactivated for 72h. Single values from n=4 independent replicates are shown. Bars represent mean ± SD. **(g)** iSLK.219 were analyzed by RTqPCR for the indicated viral targets. Single values from n=3 independent replicates are shown. Bars represent mean ± SD. **(h)** Immunoblot for the indicated viral proteins, actin as a loading control. Representative, cropped immunoblots are shown, uncropped blots are shown in Supplementary Fig.6. The experiment was done two independent times. **(i)** Immunoblot analysis for the indicated proteins using cell lysate whose luciferase activity was measured in Fig. 6b. TUBG1 as loading control. Representative, cropped immunoblots are shown, uncropped blots are shown in Supplementary Fig.6. **(j)** ChIP using the indicated antibodies followed by RTqPCR for the indicated regions upstream of the ORF50 promoter in latent iSLK.219 transduced with PROX1 WT-expressing lentivirus. Single values from n=2 independent replicates are shown. Bars represent mean ± SD. P values in panels (c), (e), (f) were calculated using ordinary one way-anova followed by Dunn´s correction for multiple comparisons *: $p<0.033$; **: $p<0.02$, ***: $p<0.001$. Exact p values are shown in Supplementary Table3.

a Fig.2b (upper panel)

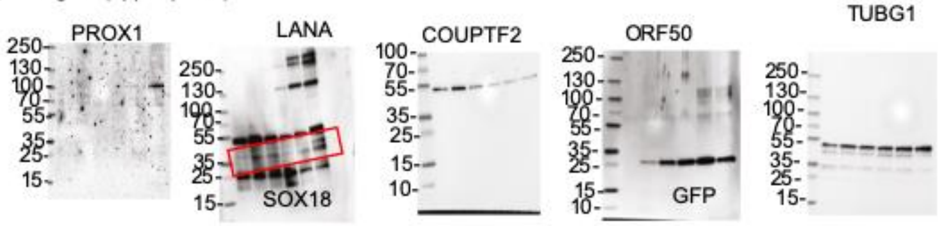


b Fig.2b (lower panel)

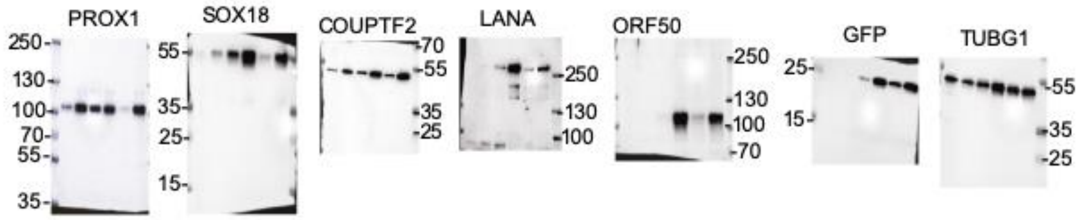


c Fig.1c

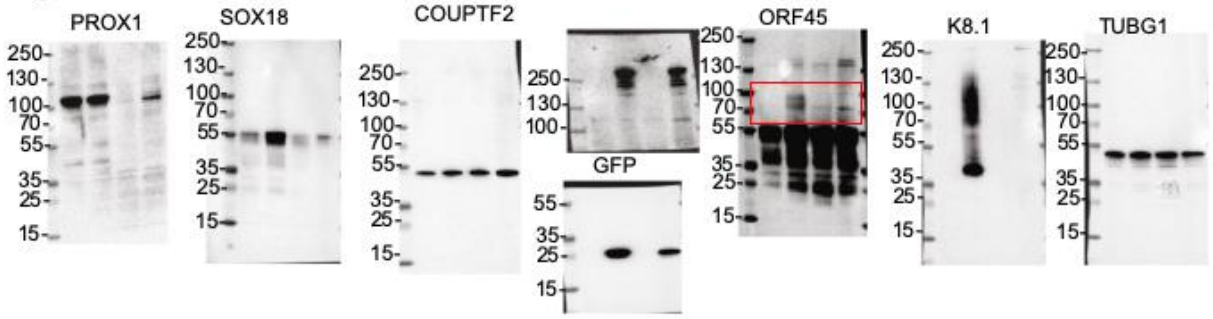


d Fig.3b

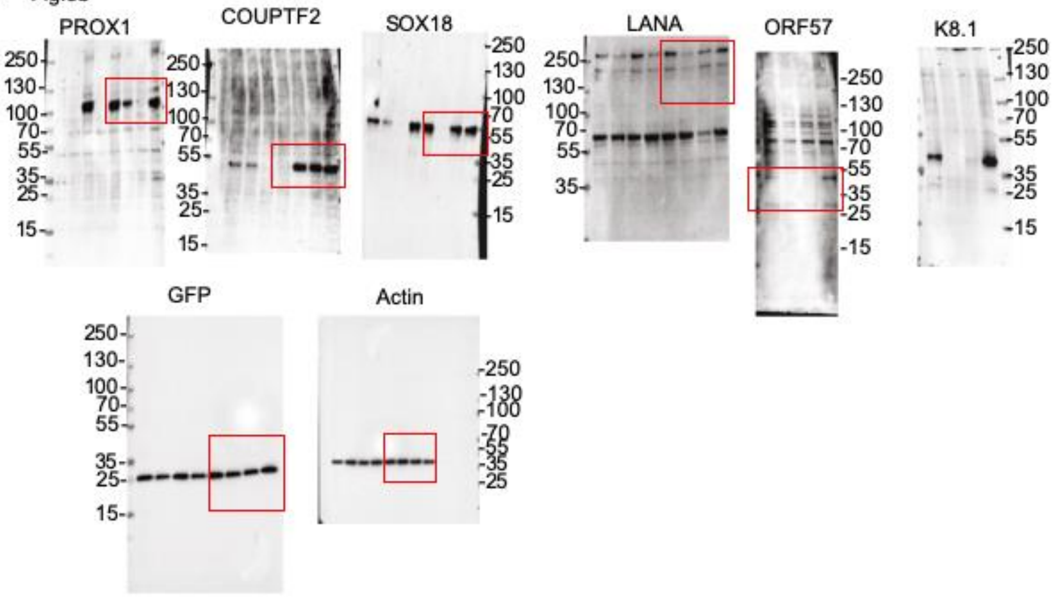


e Fig.6c

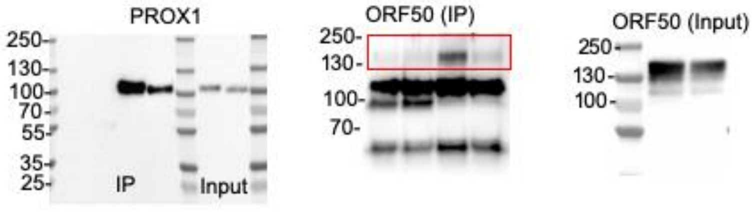

f Supplementary Fig. 2b

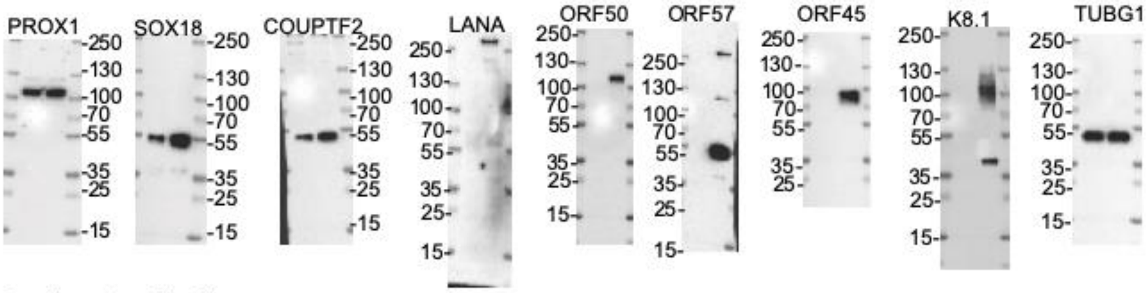


g Supplementary Fig. 3b

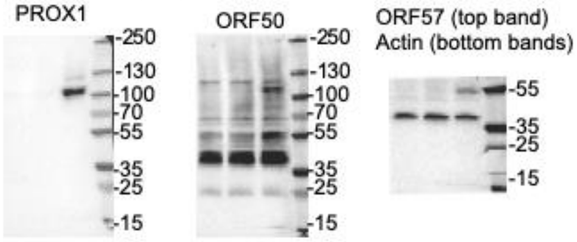


h Supplementary Fig. 3e

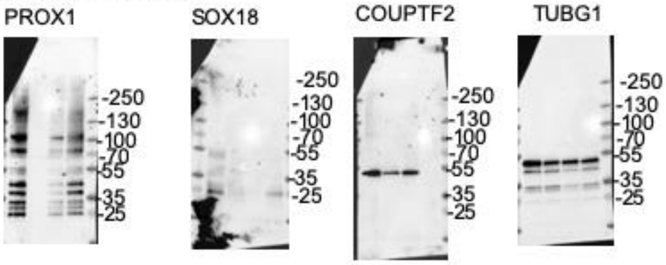


i Supplementary Fig. 4c

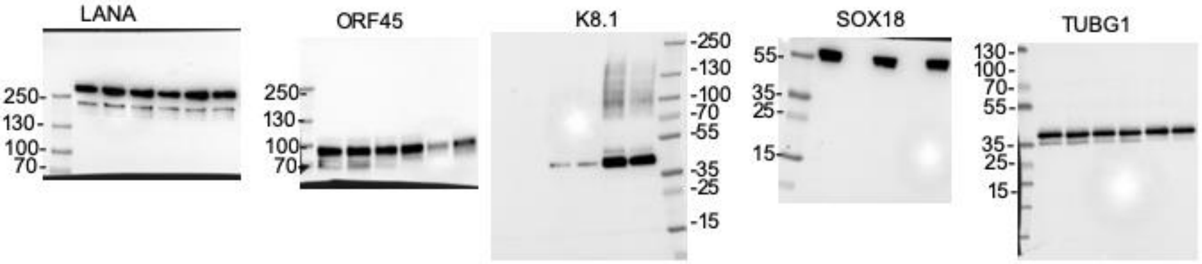


j Supplementary Fig. 4e

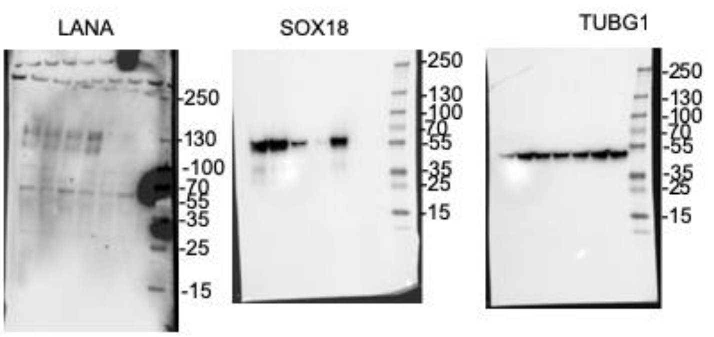


k Supplementary Fig. 4f

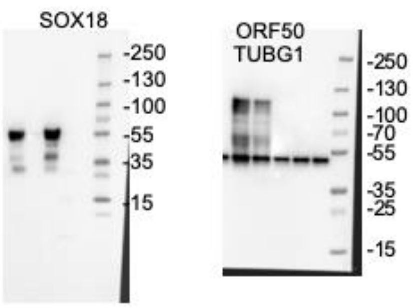

l Supplementary Fig. 5d

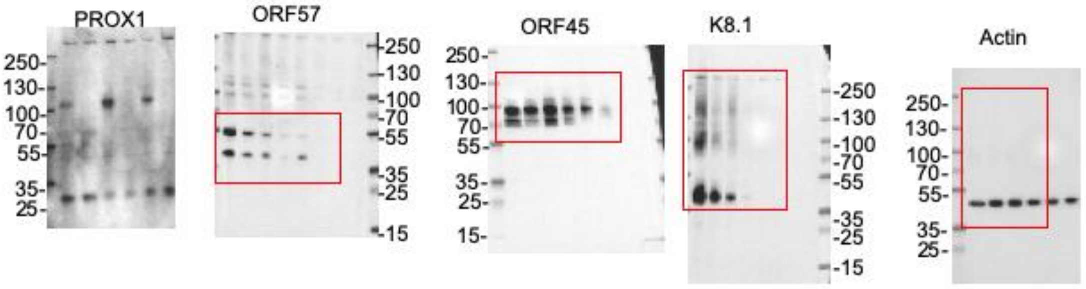


**m** Supplementary Fig. 5h

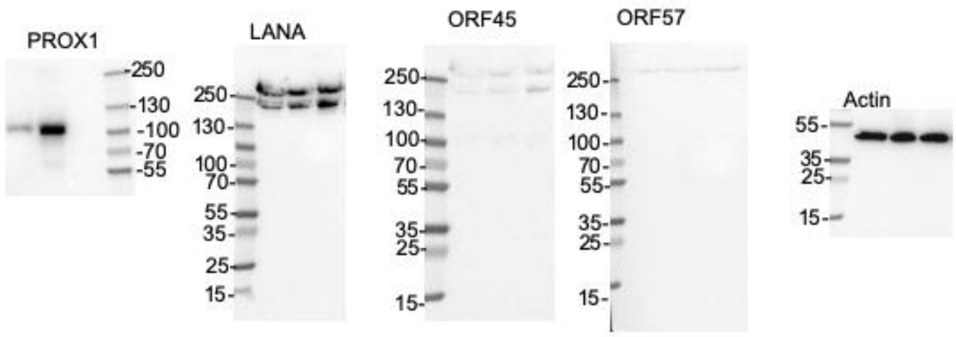


**n** Supplementary Fig. 5i

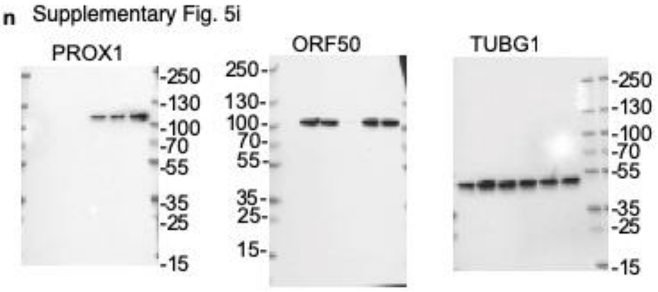


**Supplementary Fig.6.** uncropped immunoblot membranes for the indicated figures. The target protein is indicated on top of each blot, the molecular weight is shown on the side. The red square indicates the portion of the blot presented in the corresponding Figure.

**Supplementary table 1 and Supplementary table 2**: list of differentially regulated genes in KLEC and iSLK.219 treated as indicated.

**Supplementary Table 3:** Exact p values for the indicated panels.
